# Distinct regions of *H. pylori’s* bactofilin CcmA regulate protein-protein interactions to control helical cell shape

**DOI:** 10.1101/2022.04.05.487110

**Authors:** Sophie R Sichel, Benjamin P Bratton, Nina R Salama

## Abstract

The helical shape of *H. pylori* cells promotes robust stomach colonization, however, how the helical shape of *H. pylori* cells is determined is unresolved. Previous work identified helical-cell-shape-promoting protein complexes containing a peptidoglycan-hydrolase (Csd1), a peptidoglycan precursor synthesis enzyme (MurF), a non-enzymatic homologue of Csd1 (Csd2), non-enzymatic transmembrane proteins (Csd5 and Csd7), and a bactofilin (CcmA). Bactofilins are highly conserved, spontaneously polymerizing cytoskeletal bacterial proteins. We sought to understand CcmA’s function in generating the helical shape of *H. pylori* cells. Using CcmA deletion analysis, *in vitro* polymerization, and *in vivo* co-immunoprecipitation experiments we identified that the bactofilin domain and N-terminal region of CcmA are required for helical cell shape and the bactofilin domain of CcmA is sufficient for polymerization and interactions with Csd5 and Csd7. We also found that CcmA’s N-terminal region inhibits interaction with Csd7. Deleting the N-terminal region of CcmA increases CcmA-Csd7 interactions and destabilizes the peptidoglycan-hydrolase Csd1. Using super-resolution microscopy, we found that Csd5 recruits CcmA to the cell envelope and promotes CcmA enrichment at the major helical axis. Thus, CcmA helps organize cell-shape-determining proteins and peptidoglycan synthesis machinery to coordinate cell wall modification and synthesis, promoting the curvature required to build a helical cell.

## Introduction

*Helicobacter pylori* is a helical-shaped, Gram-negative bacteria that infects more than half of the world’s population (Hooi *et al*, 2017). Chronic infection with *H. pylori* can cause gastric ulcers and cancers; approximately 80% of gastric cancer cases are attributed to *H. pylori* infection (de Martel *et al*, 2020). The helical shape of *H. pylori* cells helps establish colonization of the stomach and promotes immunopathology during infection (Martinez *et al*, 2018; Montecucco & Rappuoli, 2001; Sycuro *et al*, 2010; Sycuro *et al*, 2012; Yang *et al*, 2019). Multiple proteins have been identified that are essential for the helical shape of *H. pylori* cells, most of which influence the shape and composition of the peptidoglycan (PG) sacculus directly or indirectly (Sycuro *et al*., 2010; Sycuro *et al*, 2013; Sycuro *et al*., 2012; Yang *et al*., 2019).

Several cell-shape-determining (Csd) proteins form membrane-spanning protein complexes. Csd5 is a transmembrane protein that binds directly to PG in the periplasm with its SH3 domain and binds to CcmA in the cytoplasm. Csd5 also interacts with ATP synthase and MurF, an essential PG precursor enzyme that catalyzes the final cytoplasmic PG biosynthesis step, independent of interactions with CcmA (Blair *et al*, 2018; Hrast *et al*, 2013). Another complex is formed by Csd7, Csd1, and Csd2. Csd7 is a transmembrane protein that stabilizes the PG-hydrolase Csd1, and Csd2, a catalytically inactive homologue of Csd1. Weak interactions between Csd7 and CcmA have been detected by co-immunoprecipitation experiments that include cross-linking reagents followed by mass spectrometry, but whether CcmA, Csd7, and Csd5 are all part of one protein complex is unclear (Yang *et al*., 2019). CcmA and probes that report on PG synthesis localize to the major helical axis of *H. pylori* cells (regions with a Gaussian curvature value of 5 ±1 µm^-2^ on a helical cell) indicating that CcmA is in the same region of the cell where increased levels of PG synthesis occur (Taylor *et al*, 2020).

CcmA is the only bactofilin encoded in the *H. pylori* genome. Bactofilins, a class of highly conserved, spontaneously-polymerizing, cytoskeletal proteins in bacteria, perform a variety of functions (Bulyha *et al*, 2013; El Andari *et al*, 2015; Lin *et al*, 2017; Osorio-Valeriano *et al*, 2019) including modulating cell shape in multiple different bacterial species (Brockett *et al*, 2021; Hay *et al*, 1999; Jackson *et al*, 2018; Koch *et al*, 2011; Sycuro *et al*., 2010) and morphogenesis of cell appendages (Billini *et al*, 2019; Caccamo *et al*, 2020; Kuhn *et al*, 2010), both of which involve modification or synthesis of PG. But how a cytoplasmic bactofilin influences the PG cell wall in the periplasm is unclear. Bactofilins are identified by the presence of a bactofilin domain (DUF583) which forms a β-helix and is flanked by unstructured terminal regions. The function of the terminal regions and bactofilin domain of bactofilins are debated. It has been proposed that the terminal regions may be involved in modulating polymerization, binding to membranes and/or interacting with other proteins (Deng *et al*, 2019; Kuhn *et al*., 2010; Vasa *et al*, 2015)

Here, we performed structure-function analysis to determine how the bactofilin domain and the terminal regions of CcmA contribute to generating the helical shape of *H. pylori* cells by evaluating cell morphology, CcmA polymerization and filament bundling, and interaction with known Csd protein interaction partners. Additionally, we identified how protein binding partners impact CcmA localization. We found that polymer bundling and interactions between CcmA and other Csd proteins are responsible for CcmA’s ability to pattern the helical shape of *H. pylori* cells and identified that interactions between CcmA and its binding partners modulate CcmA localization and modulate protein stability of the PG-hydrolase Csd1. Together these findings allow us to establish a new model for how *H. pylori*’s helical cell shape is generated. We clarify that CcmA participates in two separate helical-cell-shape-generating protein complexes. Additionally, these studies reveal that bactofilins can influence PG sacculus structure by modulating the protein stability of a PG hydrolase.

## Results

### CcmA is composed of a bactofilin domain and two short terminal regions

We identified the functional regions of CcmA by aligning the amino acid sequence of CcmA with BacA, a well-studied bactofilin from *C. crescentus* involved in stalk morphogenesis, using Swiss-Model (Waterhouse *et al*, 2018) and found that CcmA is composed of a bactofilin domain (DUF 583) surrounded by two short terminal regions (Figure 1A). The N-terminal region is composed of amino acids 1-17, the bactofilin domain is composed of amino acids 18-118, and the C-terminal region is composed of amino acids 119-136. Deng and colleagues proposed that bactofilins may contain a short membrane binding motif within their N-terminal regions and in CcmA that motif spans amino acids 3-12 (Deng *et al*., 2019). After identifying the boundaries between the bactofilin domain and the terminal regions of CcmA we used RoseTTAFold (Baek *et al*, 2021), a protein structure prediction service that uses deep learning-based modeling methods, to predict the structure of CcmA. RoseTTAFold generated five models of CcmA. In all five, the bactofilin domain forms a three-sided, right-handed β-helix with a left-handed twist (Figure 1B), the N-terminal region is unstructured, and part of the C-terminal region forms a short ɑ-helix and caps the end of the bactofilin domain (Figure 1C). In four of the five models, the first several amino acids of the N-terminal region interact with the bactofilin domain suggesting that the N-terminal region could potentially block a surface of the bactofilin domain.

**Figure 1.**
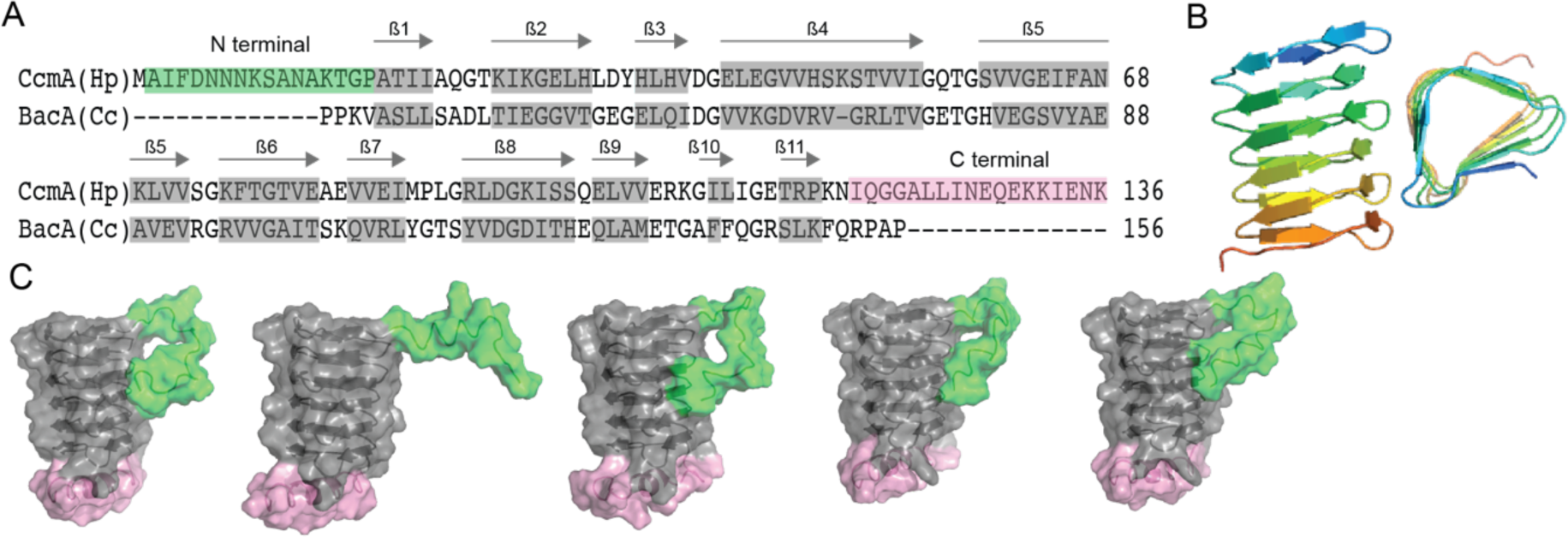
Predicted structure of CcmA. (A) Sequence alignment between BacA from *C. crescentus* and CcmA from *H. pylori* generated with SwissModel, gray boxes indicate β-strands formed in the bactofilin domain, the green box indicates the N-terminal region of CcmA and the pink box indicates the C-terminal region of CcmA. (B) Model of the bactofilin domain of CcmA generated by Robetta which forms a right-handed, 3-sided, triangular β-helix. (C) Five models of CcmA generated in RoseTTAFold displaying the bactofilin domain (gray), N-terminal region (green), and C-terminal region (pink).

### The N-terminal region and bactofilin domain are together necessary and sufficient to pattern the helical shape of *H. pylori* cells

To identify the role of each domain/region of CcmA we generated *H. pylori* strains that have truncated versions of *ccmA* at the native c*cmA* locus. We generated five strains, one strain missing the N-terminal region (Δ*NT*, missing amino acids 2-17), another strain missing the proposed membrane binding motif within the N-terminus (Δ*MM*, missing amino acids 3-12), a third strain lacking the five amino acids at the end of the N-terminus (Δ*13-17*, lacking amino acids 13-17), a fourth strain missing the C-terminal region (Δ*CT*, missing amino acids 119-136), and lastly, a strain expressing only the bactofilin domain lacking both the N-terminal region and C-terminal region (BD, expressing amino acids 18-118) (Figure 2A). The open reading frames (ORF) of the end of *csd1* and the beginning of *ccmA* overlap by 82 nucleotides (Sycuro *et al*., 2010), thus to avoid impacting the function of Csd1 by removing several amino acids from the C-terminus when truncating the N-terminus of CcmA, we expressed a second copy of *csd1* at *rd*xA, a locus commonly used for complementation (Smeets *et al*, 2000). In previous work, we confirmed that both CcmA and Csd1 can be individually deleted and complemented back without impacting the expression of the other (Sycuro *et al*., 2010; Yang *et al*., 2019).

**Figure 2.**
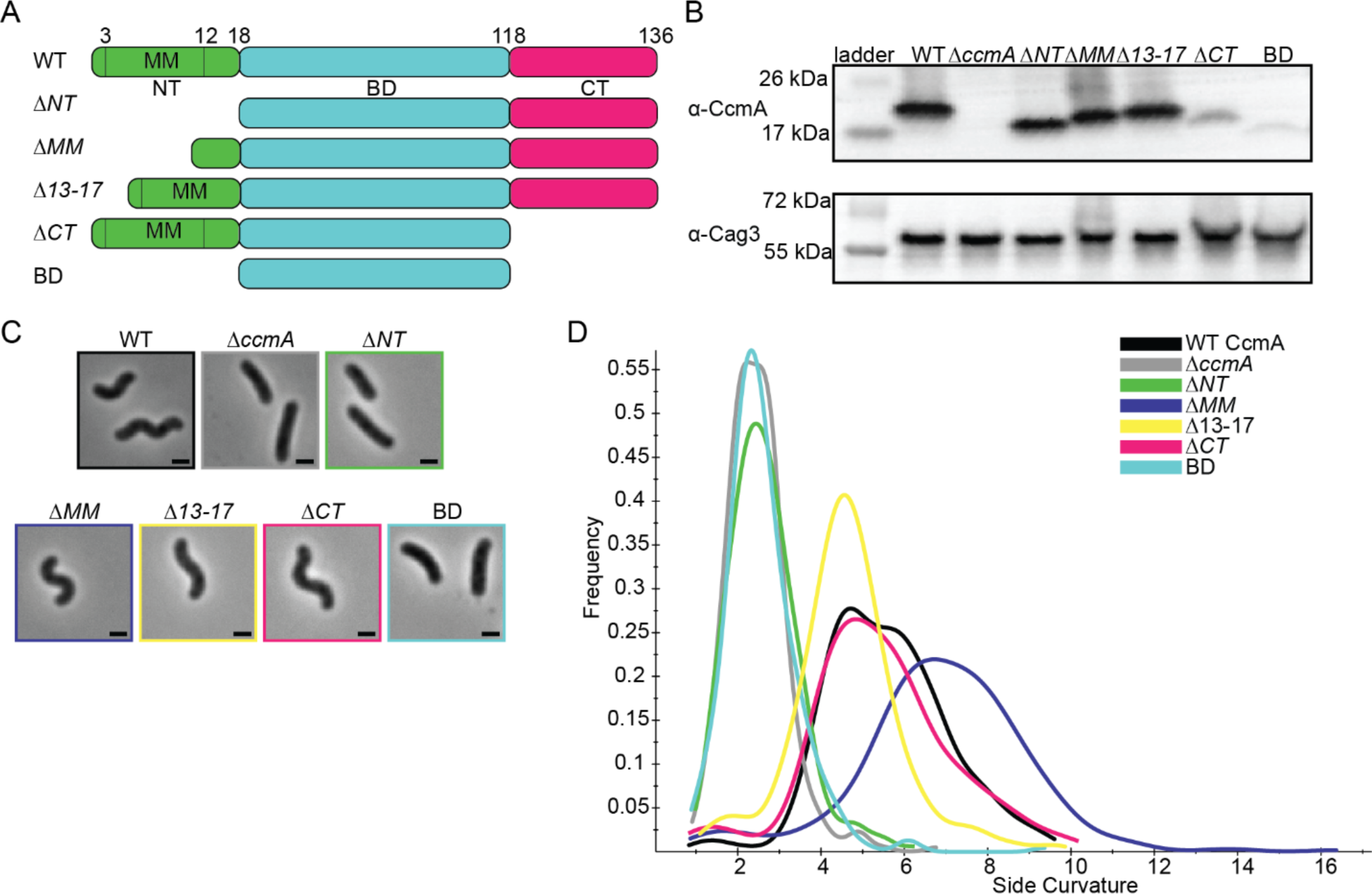
*H. pylori* cells expressing CcmA truncation mutants have different cell shapes. (A) Schematic of CcmA and truncation mutants. The N-terminal region (NT, green), bactofilin domain (BD, cyan), and C-terminal region (CT, pink) are indicated as well as the putative membrane binding motif (MM). (B) Western blot to detect CcmA in whole cell lysates of *H. pylori* strains expressing truncated versions of CcmA probed with ɑ-CcmA polyclonal antibody, and ɑ-Cag3 polyclonal antibody as a loading control. (C) Phase contrast images of *H. pylori* strains expressing truncated versions of CcmA diagramed in panel A, scalebars = 1 µm. (D) Histogram displaying the side curvature of *H. pylori* strains expressing truncated versions of CcmA (WT n= 476 cells, Δ*ccmA* n= 397 cells, Δ*N* n=275 cells, Δ*MM* n= 391 cells, Δ13-17 n=361 cells, Δ*CT* n= 263 cells, BD n=315 cells). Data are representative of two independent biological replicates. Strains used: SSH51A, SSH53A, SSH55A, SSH54A, SSH67A, SSH56B, and SSH57A.

All strains expressed the truncated versions of CcmA measured by Western blotting with an ɑ-CcmA polyclonal antibody, however, in strains lacking the C-terminus (Δ*CT* and BD) we saw much lower signal on the blot (Figure 2B). In follow-up experiments we determined that the ɑ-CcmA polyclonal antibody fails to recognize versions of CcmA lacking the C-terminus (Supplemental Figure 1). We ran a Western blot probed with our polyclonal ɑ-CcmA antibody on purified 6-his WT CcmA and CcmA truncation mutants (Supplemental Figure 2). Although equal concentrations of each purified protein were loaded, the polyclonal ɑ-CcmA antibody poorly recognized versions of CcmA lacking the C-terminal region (Δ*CT* and BD), suggesting that the Δ*CT* and BD proteins are likely expressed in *H. pylori* cells at similar levels to other versions of CcmA but are not robustly recognized by the ɑ-CcmA polyclonal antibody.

Next, after confirming that all truncated versions of CcmA were expressed, we assessed the shapes of the cells and found that strains lacking the N-terminal region (Δ*NT* and BD) were indistinguishable from Δ*ccmA* cells (Figure 2 C-D), while cells lacking the C-terminal region of *ccmA* resembled WT cells. Cells that lacked either the putative membrane binding motif (Δ*MM*) or the last five amino acids of the N-terminal region (Δ*13-17*) were helical but had slightly different side curvature values than the WT population, suggesting that the N-terminal region of CcmA may have multiple functional regions within it.

### Circular dichroism spectroscopy reveals that CcmA truncation mutants maintain β-strand secondary structure

Circular dichroism (CD) spectroscopy can be used to identify the secondary structure and folding of proteins. Different structural elements have characteristic CD spectra and proteins that consist of β-strands have negative peaks at 216 nm (Greenfield, 2006). We sought to identify whether CcmA is composed of β-strands and confirm that our truncated versions of CcmA were still folded correctly, especially considering that in *H. pylori* strains lacking the N-terminal region of CcmA, cells resembled Δ*ccmA* mutants. We recombinantly expressed and purified 6-his CcmA (Supplemental Figure 2) and collected circular dichroism spectra from 200-260 nm at fixed temperatures to identify the secondary structure and measure its thermal stability.

CD spectra of WT CcmA and the truncation mutants Δ*NT*, Δ*CT*, and BD at 20 °C all look very similar; they have the characteristic negative peak at 216 nm indicating β-strand secondary structure (Figure 3A). Together, these data combined with the modeling by RoseTTAFold in Figure 1C suggest that CcmA’s bactofilin domain very likely forms a β-helix. In contrast, we also ran CD scans on WT CcmA at 95 °C and two CcmA point mutants that our group previously studied, I55A and L110S at 20 °C (Taylor *et al*., 2020). The CD spectra of both mutants (I55A and L110S) at 20 °C resemble spectra of melted WT CcmA at 95 °C (Figure 3B), suggesting that these proteins are not folded. Thus, while CcmA Δ*NT*, BD, I55A, and L110S all have the same cell shape phenotype when expressed in *H. pylori* - loss of helical cell shape - the mechanism differs. In CcmA I55A and L110S, CcmA is not folded and nonfunctional, while in CcmA Δ*NT* and BD, the protein is folded, but loss of the N-terminus causes a loss of helical cell shape.

**Figure 3.**
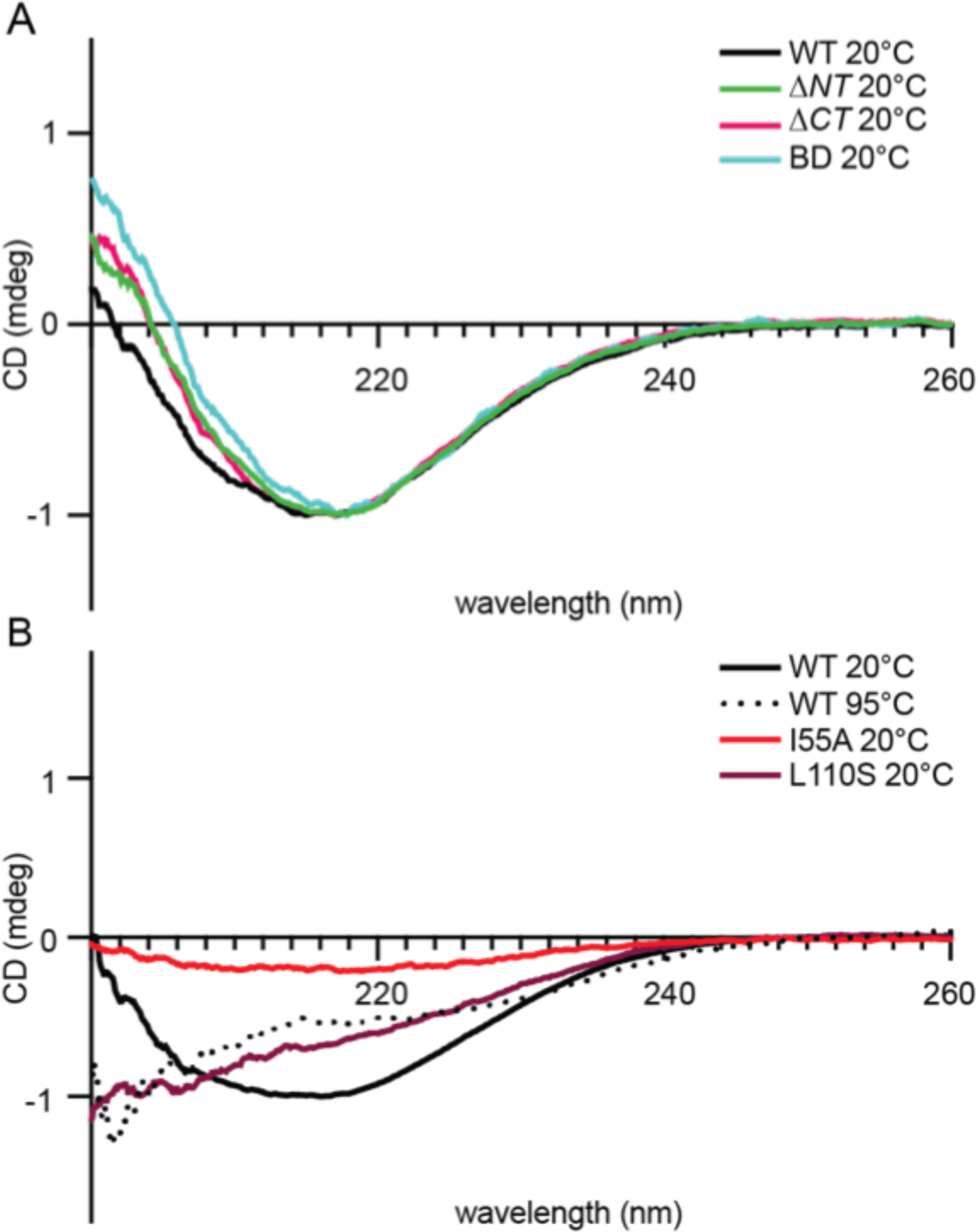
Circular dichroism spectra of purified 6-his CcmA suggests CcmA is composed of predominantly β-sheets. (A) Circular dichroism spectra of WT and truncated versions of CcmA at room temperature (20 °C), WT (black), Δ*NT* (green), Δ*CT* (pink), and BD (cyan). (B) Circular dichroism spectra of WT CcmA at room temperature (20 °C, black), melted (95 °C, dotted line), and two point mutants (I55A, red and L110S, burgundy).

### The bactofilin domain alone is capable of polymerization; the terminal regions of CcmA modulate lateral interactions between CcmA filaments

To identify how each domain/region of CcmA contributes to *in vitro* polymerization we expressed and purified truncated versions of 6-his CcmA and assessed polymerization by negative stain and transmission electron microscopy (TEM). As previously reported (Holtrup *et al*, 2019; Taylor *et al*., 2020), WT CcmA polymerizes into small filaments that measure on average 3.04 nm (SD= 0.79, n=18) in width. The filaments wrap together to form bundles of approximately 21.34 nm (SD= 4.82, n=24) wide, and form lattice structures (Figure 4 A, G). Δ*NT* formed filaments and bundles which were more homogenous, straighter, and thinner than WT and appear to only contain 2-4 filaments each. Δ*NT* failed to form lattice structures (Figure 4B). Δ*MM* resembled WT CcmA, forming filaments and bundles of approximately the same width as WT and formed lattices (Figure 4C, G). Δ*13-17* resembled Δ*NT* (Figure 4D, G), suggesting that amino acids 13-17 are responsible for filament bundling properties of Δ*NT*. Δ*CT* formed bundles that were wider than WT CcmA but failed to form lattices (Figure 4E, G). Lastly, BD formed straight bundles that were narrower than those found in WT samples, and failed to form lattices (Figure 4F, G). These data suggest that the bactofilin domain is sufficient for polymerization of CcmA while the N- and C-terminal regions of CcmA regulate inter-polymer interactions that impact the width of bundles by modulating the number of filaments that form a bundle, and the ability of CcmA to form lattices.

**Figure 4.**
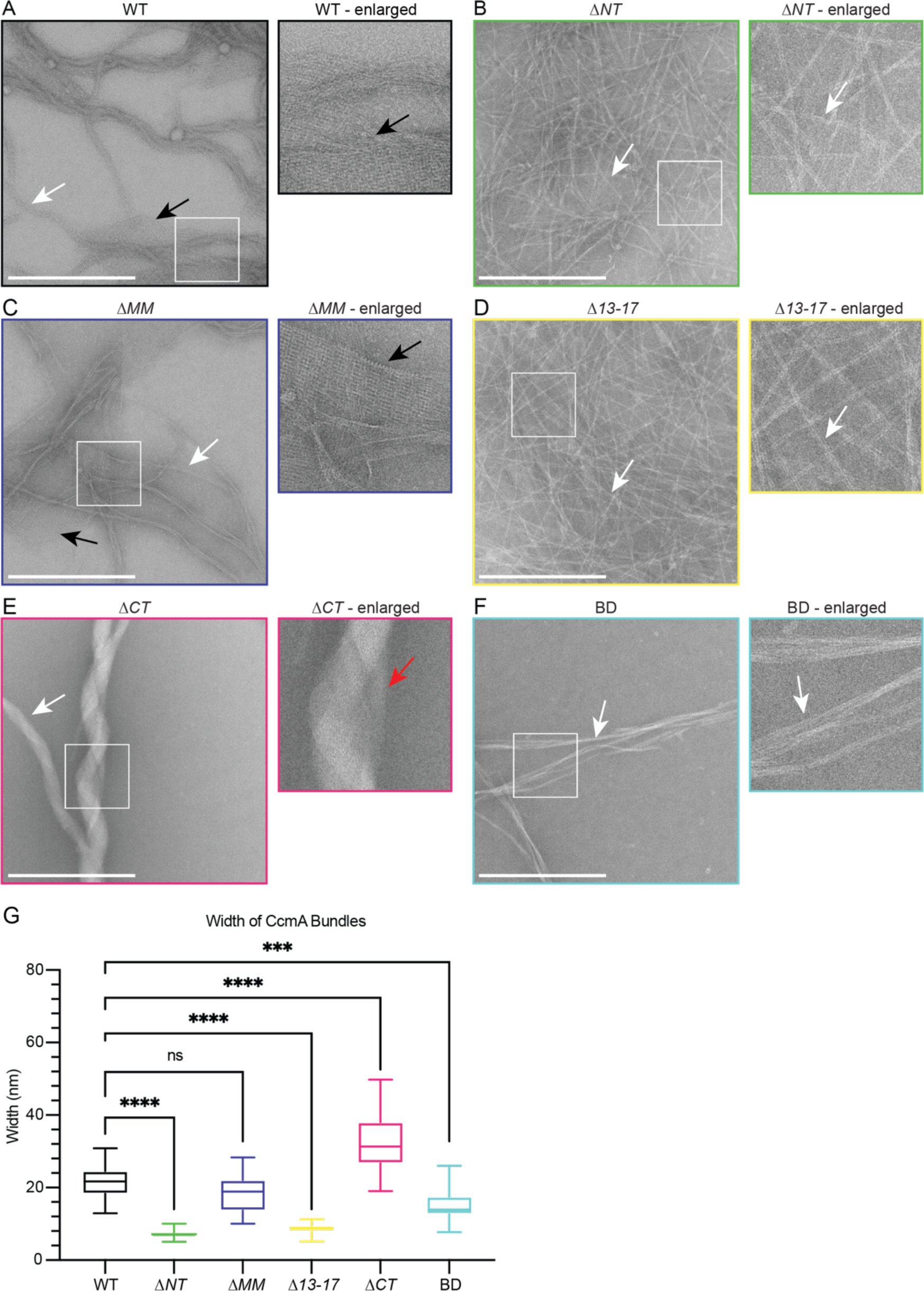
The bactofilin domain of CcmA is sufficient for *in vitro* polymerization while the terminal regions promote lateral polymer interactions. (A-F) TEM of negative stained, purified CcmA, scalebars = 500 nm. White boxes indicate region of micrograph that is enlarged in panel to the right of original. White arrows indicate bundles of CcmA, black arrows indicate lattice structures, red arrow indicates two bundles wrapping around each other. Data are representative of two independent biological replicates. (G) Box and whiskers plot displaying width of purified CcmA bundles and filaments. Median, min, max, 25^th^ percentile, and 75^th^ percentile are displayed. One-way ANOVA, Dunnett’s multiple comparisons test. ns P>0.5, *** P≤0.001, **** P ≤0.0001. WT n=24, Δ*NT* n=29, Δ*MM* n=25, Δ*13-17* n=31, Δ*CT* n=24, BD n=32.

### CcmA’s bactofilin domain mediates interactions with Csd5 and Csd7, and the N-terminal region of CcmA inhibits interactions with Csd7

CcmA was previously shown to interact with two Csd proteins, Csd5 and Csd7, by co-immunoprecipitation experiments. Csd5-CcmA interactions were readily detected by co-immunoprecipitation followed by mass spectrometry and by Western blotting. However, Csd7-CcmA interactions could only be detected when a cross-linking reagent was included in co-immunoprecipitation experiments and mass spectrometry was used to detect CcmA, suggesting that Csd7-CcmA interactions are weak or potentially transient (Blair *et al*., 2018; Yang *et al*., 2019). To probe how each region of CcmA contributes to interactions with Csd5 and Csd7, we performed co-immunoprecipitation experiments followed by Western blotting. Strains were generated that expressed truncated versions of *ccmA* at the native *ccmA* locus, *csd1* at the *rdxA* locus as described earlier, and either *csd5-2X-FLAG* or *csd7-3X-FLAG* at their native loci. We immunoprecipitated either Csd5-FLAG or Csd7-FLAG with ɑ-FLAG agarose beads, then performed Western blotting and probed with the ɑ-CcmA polyclonal antibody to identify whether the truncated versions of CcmA could still co-purify. When we immunoprecipitated Csd5-FLAG, all mutant versions of CcmA were detected by Western blotting of the IP fractions, except for the Δ*CT* mutant (Figure 5A), though it is likely that this mutant can still interact with Csd5-FLAG. Earlier we identified that the C-terminal region of CcmA is required for robust detection by our ɑ-CcmA polyclonal antibody (Supplemental Figure 1) and here, the BD mutant interacts with Csd5. Additionally, we tagged WT CcmA, Δ*CT* and BD with a HaloTag in *H. pylori* at the native *ccmA* locus and performed a co-immunoprecipitation experiment to detect interactions with Csd5-FLAG (Supplemental Figure 3). We found that both Δ*CT*-HaloTag and BD-HaloTag were expressed at comparable levels as CcmA-HaloTag and could be pulled down by Csd5-FLAG. These three pieces of evidence lead us to conclude that the Δ*CT* mutant can interact with Csd5, suggesting that only the bactofilin domain is required for interactions between CcmA and Csd5-FLAG.

**Figure 5.**
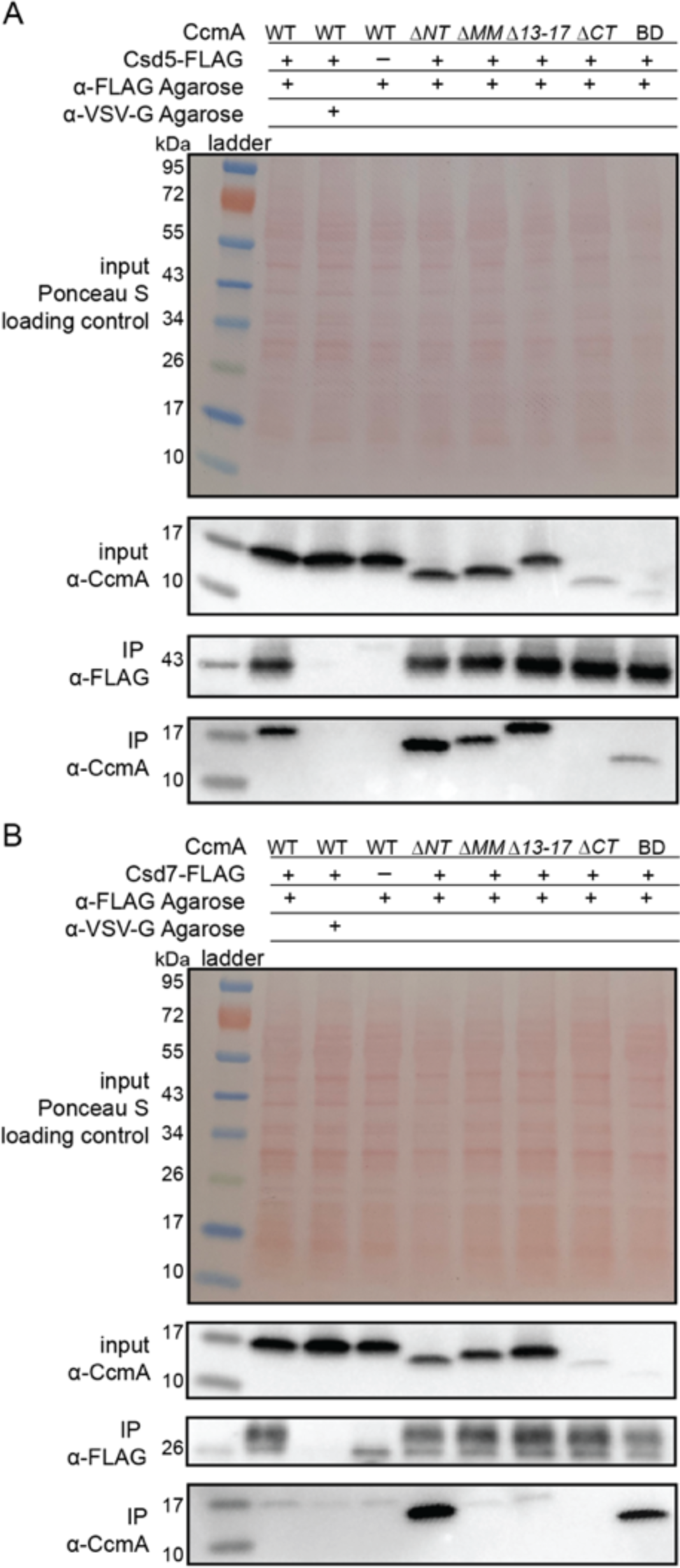
The bactofilin domain interacts with Csd5 and Csd7, and the N-terminal region inhibits Csd7 binding to CcmA. Co-immunoprecipitation experiments to probe Csd5-2x-FLAG-CcmA interactions (A) and Csd7-3x-FLAG-CcmA interactions (B) in *H. pylori* cells are shown. Top row: Ponceau S staining of input fractions. Second row: Western blot probed with ɑ-CcmA polyclonal antibody of input fractions. Third row: Western blot probed with ɑ-FLAG monoclonal antibody of immunoprecipitation (IP) fractions. Bottom row: Western blot probed with ɑ-CcmA polyclonal antibody of IP fractions. Data shown are representative of data from three independent biological replicates. Strains used: KHB157, LSH100, SSH59A, SSH58A, SSH68A, SSH60A, SSH65A, DCY71, SSH79A, SSH78A, SSH82, SSH80A, and SSH81B.

When we immunoprecipitated Csd7-FLAG we found that, as previously reported (Yang *et al*., 2019), WT CcmA was not able to be detected in the IP fractions by Western blotting above background levels. However, versions of CcmA lacking the N-terminal region were robustly co-purified by Csd7 (Figure 5B) suggesting that the N-terminal region inhibits WT CcmA from being co-immunoprecipitated by Csd7-FLAG. Together these data suggest that the bactofilin domain of CcmA mediates interaction with Csd5 and Csd7, the N-terminal region of CcmA inhibits interaction with Csd7, and versions of CcmA lacking the C-terminal region are expressed at comparable levels to WT CcmA although they are not recognized as robustly by the ɑ-CcmA polyclonal antibody.

### Increased CcmA-Csd7 interactions preclude Csd1-Csd7 interactions, resulting in decreased Csd1 expression

After discovering that the N-terminal region of CcmA could inhibit interactions with Csd7 and that Δ*NT* CcmA was readily co-purified by Csd7, we reasoned that Δ*NT* CcmA could be binding to Csd7 and reducing its capacity to stabilize Csd1, causing decreased levels of Csd1 expression. This hypothesis was bolstered by the finding that Δ*csd7, Δcsd1*, and Δ*NT* cells all share the same curved rod shape phenotype. To test this hypothesis, we assayed Csd1 expression by Western blotting and probing with an ɑ-Csd1 polyclonal antibody in whole cell lysates of the *H. pylori* strain lacking the N-terminal region of *ccmA*. As discussed previously, this strain express two copies of Csd1; there is an extra copy of *csd1* at *rdxA* which was added to account for potentially impacting Csd1’s function when truncating *ccmA* (the ORF of the end of *csd1* and the beginning of *ccmA* overlap by 82 nucleotides). Strikingly, we found that in the strain lacking the N-terminal region (Δ*NT*) Csd1 levels are lower than in a strain expressing WT CcmA (Figure 6 A-B). After identifying that Csd1 levels were depleted when the N-terminal region of CcmA was missing, we sought to identify if Csd7-Csd1 interactions were diminished when Csd7-CcmA interactions increased. We performed Western blotting and probed for Csd1 levels in the samples we collected during our Csd7-FLAG co-immunoprecipitation experiments (Figure 5B). We found that in *H. pylori* strains lacking the N-terminal region of CcmA, Csd7-Csd1 interactions are decreased in comparison to when WT CcmA is expressed. Suggesting that when the N-terminal region of CcmA is deleted CcmA-Csd7 interactions increase and inhibit Csd7 from stabilizing Csd1, causing decreased Csd1 levels. Our data suggest that Csd1 stability is indirectly regulated by CcmA via Csd7.

**Figure 6.**
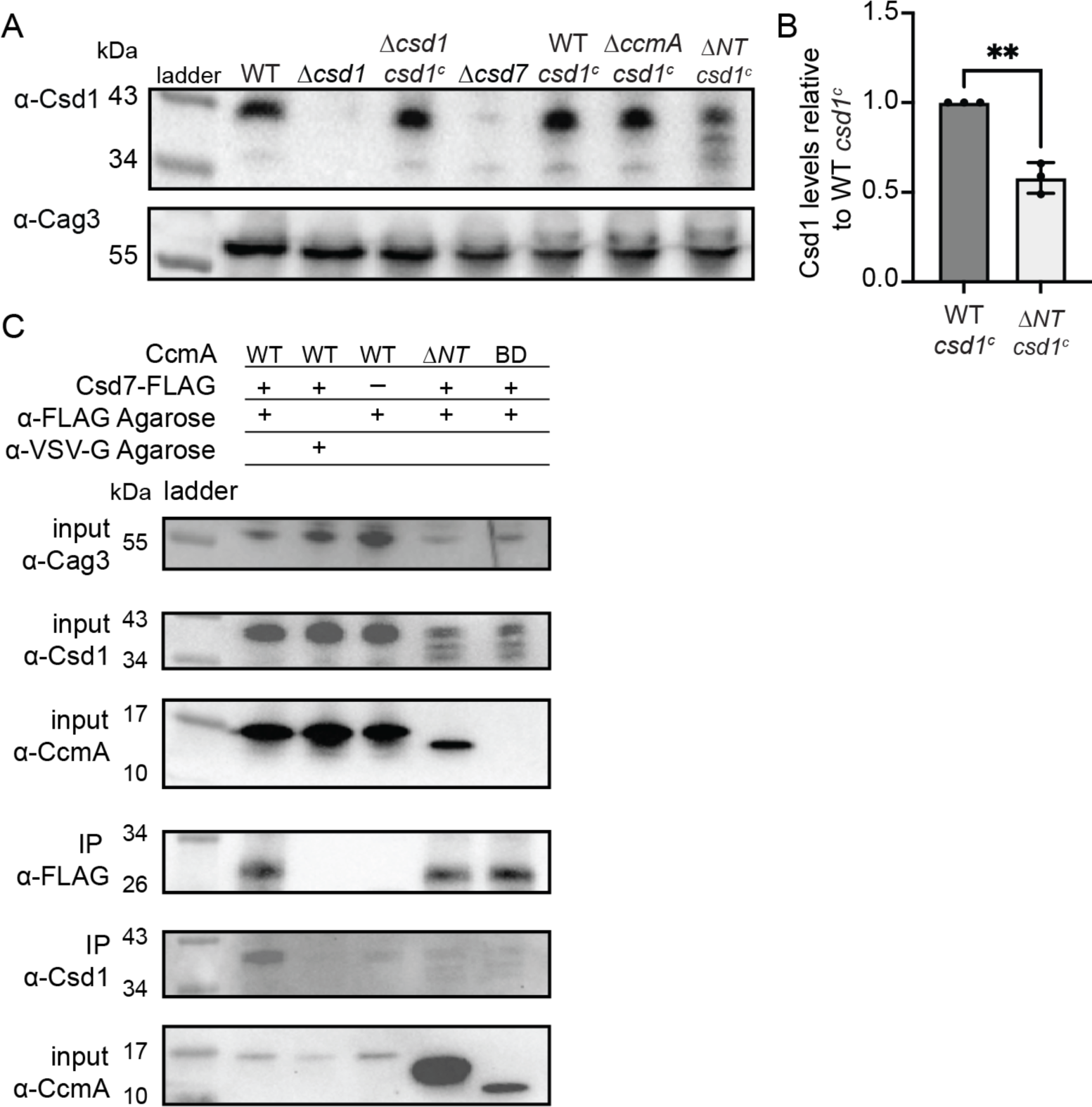
Csd1 expression is diminished when the N-terminal region of CcmA is deleted due to increased Csd7-CcmA interactions that preclude Csd7-Csd1 interactions. (A) Western blot of whole cell extracts probed with ɑ-Csd1 polyclonal antibody showing expression of Csd, ɑ-Cag3 polyclonal antibody was used as a loading control. Strains with “csd1^c^” indicate an extra copy of *csd1* was added at the *rdxA* locus. Data shown are representative of data from three independent biological replicates. (B) Quantification Csd1 levels in the ccmA Δ*N*T strain relative to the WT *csd1^c^*. Relative signal was calculated by dividing Csd1 signal by Cag3 signal for each replicate, then the data was normalized by dividing it by the value from the WT *csd1^c^* strain from the same experiment. Data are pooled from three independent biological replicates. Unpaired t test, ** P≤0.001. (C) Co-immunoprecipitation experiments to probe Csd1 levels in relation to Csd7-3x-FLAG-CcmA interactions in *H. pylori* cells are shown. Top row: Western blot of input fractions probed with ɑ-Cag3 polyclonal antibody as loading control. Second row: Western blot of input fractions probed with ɑ-Csd1 polyclonal antibody. Third row: Western blot of input fractions probed with ɑ-CcmA polyclonal antibody. Fourth row: Western blot probed with ɑ-FLAG monoclonal antibody of immunoprecipitation (IP) fractions to detect Csd7-FLAG. Fifth row: Western blot probed with ɑ-Csd1 polyclonal antibody of IP fractions. Bottom row: Western blot probed with ɑ-CcmA polyclonal antibody of IP fractions. Data shown are representative of data from two independent biological replicates. Strains used: LSH100, LSH113, LSH121, DCY26, SSH51A, SSH53A, SSH55A, DCY71, SSH79A, and SSH81B.

### CcmA requires binding partners for proper localization to the major helical axis

It has been hypothesized that bactofilins are able to interact with the cell envelope directly and sense specific curvature values. To address whether binding partners are required to direct CcmA to the cell envelope, we first engineered an *H. pylori* strain we could use to visualize CcmA. We engineered a strain with *ccmA-HaloTag* at the native *ccmA* locus and a second copy of WT *ccmA* at *rdxA*. We found that cells that expressed CcmA-HaloTag as the sole copy of CcmA were still helical but had slightly lower side curvature than WT cells; adding a second copy of *ccmA* at *rdxA* generated cells with side curvature more similar to WT cells (Supplemental Figure 4). To identify whether CcmA is capable of localizing to the cell envelope at its preferred curvature alone or whether localization is mediated by a binding partner, we deleted *csd5* and *csd7* individually and together in our *ccmA-HaloTag* strain. As an additional control we deleted *csd6* in the *ccmA-HaloTag* strain. Csd6 is a peptidoglycan carboxypeptidase that does not appear to interact with CcmA, Csd5, or Csd7 (Blair *et al*., 2018; Sycuro *et al*., 2013; Yang *et al*., 2019). Similar to Δ*csd5* mutants, Δ*csd6* mutants show a straight rod shape phenotype. We fixed and permeabilized cells, then labeled CcmA-HaloTag with a fluorescent ligand (JF-549) that binds to HaloTag and counterstained with fluorescent wheat germ agglutinin (WGA), which binds to PG, to label the cell envelope. We collected images of hundreds of cells from each population using 3D structured illumination microscopy (SIM) and used software previously described to generate 3D reconstructions of individual cells from the WGA channel and calculate the Gaussian curvature at each point on the cell surface. Then we performed analysis to identify how much CcmA-HaloTag signal was at the cell envelope versus in the center of cells, and curvature enrichment analysis to investigate whether CcmA-HaloTag is enriched or depleted at specific Gaussian curvature values (Bratton *et al*, 2019; Taylor *et al*., 2020).

In WT cells, we found that CcmA-HaloTag signal is present as foci and small arcs that colocalize with the cell envelope (white signal) and there were some foci in the center of the cell (Figure 7A top row). The CcmA-HaloTag signal closely resembled what we saw when we visualized CcmA-FLAG by immunofluorescence in previous work (Taylor *et al*., 2020). In cells lacking *csd6*, we found that CcmA-HaloTag signal is present at the cell envelope and there are some foci in the center of cells (Figure 7A second row). In cells lacking *csd7*, cells are slightly curved and CcmA-HaloTag signal colocalizes with the cell envelope, however there are also some foci in the center of the cell (Figure 7A third row). In cells lacking *csd5*, there appears to be more CcmA-HaloTag signal in the center of cells than associated with the cell envelope (Figure 7A fourth row). In cells lacking both *csd5* and *csd7*, we found that the cell shape resembles a Δ*csd7* mutant, however, there appears to be more CcmA-HaloTag signal in the center of the cells than in the Δ*csd7* mutant, similar to the Δ*csd5* mutant (Figure 7A bottom row).

**Figure 7.**
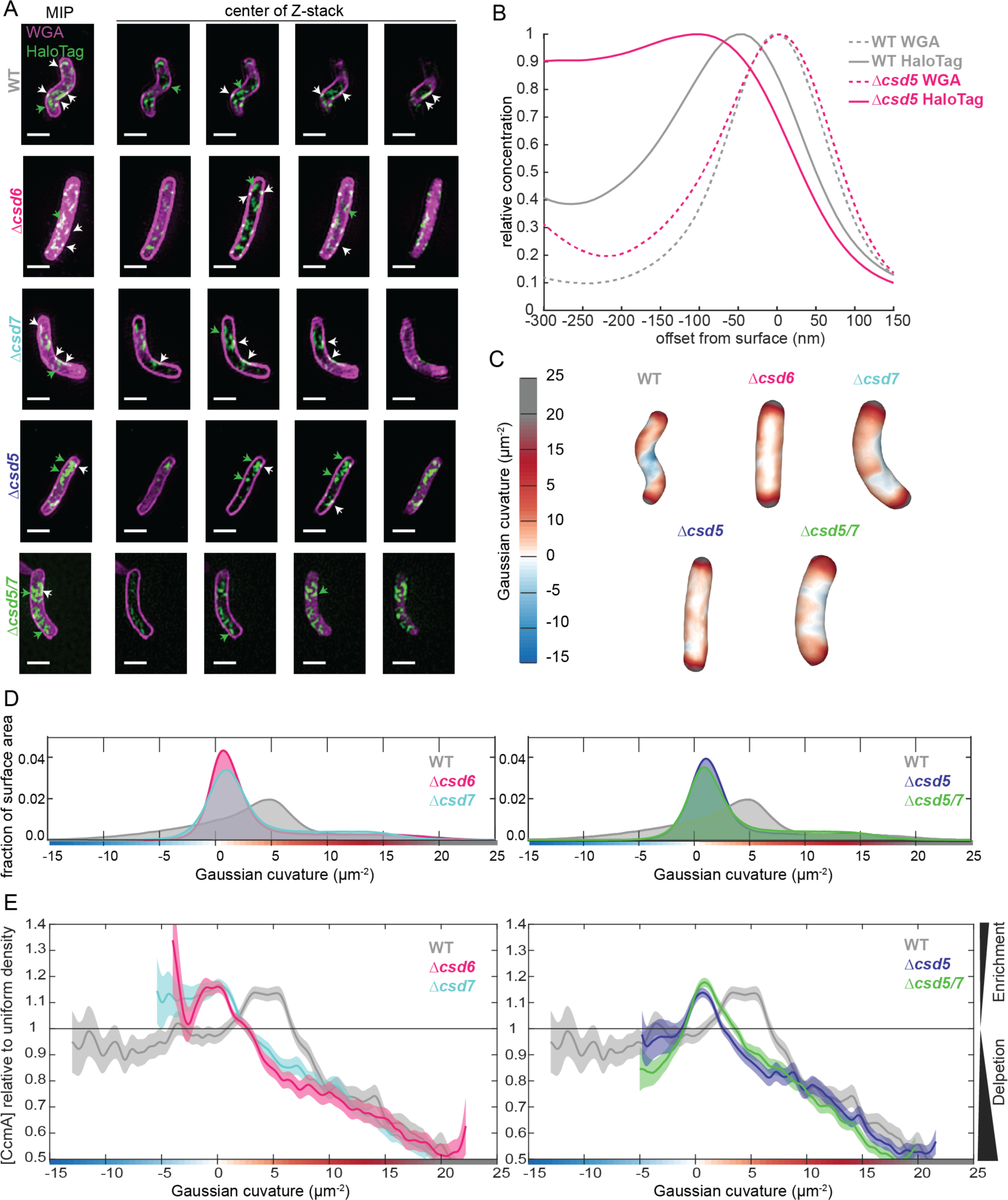
Csd5 is required for CcmA-HaloTag to localize to the cell envelope at the major helical axis. (A) Maximum intensity projections (MIP) and frames from Z-stacks of *H. pylori* cells labeled with WGA to label the cell wall (magenta) and JF549 ligand to label CcmA-HaloTag (green), pixels are white where the two signals co-localize. Green arrows indicated cytoplasmic CcmA-HaloTag. White arrows indicate CcmA-HaloTag signal that colocalizes with the cell envelope. Scalebars = 1 µm. (B) Representative plots displaying the relative concentration of HaloTag-CcmA and WGA signal from one WT and one Δ*csd5* cell at the cell surface (X=0) and at computationally generated cell surfaces that are inside (X<0) and outside (X>0) of the cell surface. (C) Representative *H. pylori* cells from each population with Gaussian curvature values mapped on the cell surface. (D-E) Top, histogram of Gaussian curvature of cell surfaces of each population. Bottom, surface Gaussian curvature enrichment of relative concentration of CcmA signal in each population where 1=uniform concentration of CcmA, shaded regions indicate SEM. In panels C-F, WT n=292, Δ*csd6* n=264 cells, Δ*csd7* n=355, Δ*csd5* n=326, Δ*csd5/7* n=523. Strains used: SSH39B, SSH49A, SSH50A, JS09, and SSH70A.

To quantify how much CcmA-HaloTag signal was associated with the cell envelope as compared to the amount of CcmA-HaloTag found in the center of the cells, we developed a method to measure the amount of CcmA-HaloTag signal that colocalized with the cell surface as we computationally generated sequential cell surface shells with smaller or larger diameters (Figure 7B and Supplemental Figure 4C-D). We used the fluorescent WGA channel which labels the PG cell wall to generate the cell surface. As we computationally shrink the cell surface (X decreases), we observe the relative concentration of WGA and CcmA-HaloTag signal with respect to the position of the cell surface. In both a WT and Δ*csd5* cell the WGA signal peaks at X=0 nm where the PG cell wall is located within the periplasm. In WT cells, CcmA-HaloTag signal peaks near -50 nm from the PG cell wall and declines further to the inside of the cell. Based on cryo-electron tomography data, an approximate distance of 50 nm from the PG cell wall to the cytoplasm in *H. pylori* is expected (Asmar *et al*, 2017; Chang *et al*, 2018; Kuhn *et al*., 2010; Lu *et al*, 2022; Qin *et al*, 2017). In contrast, in a Δ*csd5* cell, the HaloTag-CcmA signal peaks near -100 nm from the cell surface and continues to be high even as the surface of interest approaches center of the cell (approximately -300 nm), indicating that more CcmA-HaloTag signal is found near the center of the cell in Δ*csd5* cells than in WT cells. When we performed the same analysis on the entire populations of cells (292 WT and 326 Δ*csd5*), we found that while the overall trends were the same as when we analyzed individual cells, there was a lot of variability in the WT cell population. The variability of the WT population arises because many WT cells have both a population of CcmA-HaloTag at the cell envelope (peak at -50 nm) and in the center of the cell that causes a broader left tail in the population level data (Supplemental figure 4C-D). In contrast, the *!1csd5* population level signal peak is broader and shifted towards the center of the cell (-80 nm). Thus, Csd5 is required for CcmA to localize to the cell envelope.

To identify where CcmA-HaloTag signal that is at the cell surface is in relation to cell curvature, we calculated the Gaussian curvature values of all cell surfaces represented in the populations. WT cells (which are helical) have a larger range of Gaussian curvature values on their cell surfaces (Figure 7 C-D) compared to all other shape mutants. Interestingly, although all shape mutants are not helical, they do display different surface Gaussian curvature distributions and Δ*csd7* and Δ*csd5/7* populations look nearly identical, suggesting that Δ*csd5/7* double mutants resemble the shape of a Δ*csd7* mutant.

In WT cells, CcmA-HaloTag is enriched at the major helical axis (Gaussian curvature value of 5 ±1 µm^-2^ (Taylor *et al*., 2020)); enrichment peaks at +3.35 µm^-2^ to +5.31 µm^-2^ Gaussian curvature and is depleted at Gaussian curvature values lower than +0.97 µm^-2^ and higher than +6.67 µm^-2^. These data are consistent previous experiments where CcmA-FLAG was detected by immunofluorescence (Taylor *et al*., 2020). In contrast, in all other strains, CcmA-HaloTag is depleted at regions of Gaussian curvature where CcmA-Halotag is enriched in WT cells (+3.35 µm^-2^ to +5.31 µm^-2^) although those Gaussian curvature values are still present in all cell-shape-mutant populations (Figure 7 E-F). In Δ*csd6* and Δ*csd7* mutants, CcmA-HaloTag is enriched at negative Gaussian curvature values. In Δ*csd6* mutants, enrichment peaks at and continues to be enriched at all values lower than +0.11 µm^-2^ Gaussian curvature (to -4 µm^-2^ Gaussian curvature). In Δ*csd7* mutants, CcmA-HaloTag enrichment peaks at +0.15 µm^-2^ and is enriched at all values below +0.15 µm^-2^ (to -5.4 µm^-2^ Gaussian curvature), (Figure 7 E). In contrast to Δ*csd6* and Δ*csd7* mutants, in Δ*csd5* mutants, CcmA-HaloTag is depleted at negative Gaussian curvature values (Figure 7 F). Interestingly, in a Δ*csd5/7* double mutant, the CcmA-HaloTag enrichment pattern mirrors the Δ*csd5* mutant enrichment pattern. In Δ*csd5* mutants CcmA-HaloTag is enriched at Gaussian curvature values from -1.18 µm^-2^ to +2.5 µm^-2^ and has a peak enrichment at +0.68 µm^-2^, CcmA-HaloTag is depleted at all other values of Gaussian curvature. In a Δ*csd5/7* double mutant CcmA-HaloTag is enriched at Gaussian curvatures of -0.99 µm^-2^ to +3.46 µm^-2^ and peaks at +0.85 µm^-2^, CcmA-HaloTag is depleted at all other values of Gaussian curvature. Suggesting that when helical cell shape is disrupted but Csd5 is present (in a Δ*csd6* or Δ*csd7* strain), CcmA localizes to regions of negative Gaussian curvature instead of the major helical axis and that Csd5, not Csd7, modulates CcmA localization.

## Discussion

The PG sacculus is what ultimately controls the shape of bacterial cells and it has been hypothesized that modification of PG crosslinking generates curvature and twist of the sacculus (Huang *et al*, 2008; Sycuro *et al*., 2010). In *H. pylori*, when the cytoplasmic bactofilin CcmA is lost, cells lose their helical shape and the PG muropeptide composition is altered (Sycuro *et al*., 2010). Interestingly, Δ*csd1*, Δ*csd2*, Δ*csd7*, and Δ*ccmA* mutants all share the same curved rod phenotype and PG muropeptide composition, suggesting that CcmA may regulate Csd1 activity, however prior to this work there was not a clear mechanism linking CcmA and Csd1 activity (Sycuro *et al*., 2010; Yang *et al*., 2019). Past work identified that CcmA interacts with the transmembrane protein Csd5 (which binds directly to PG) and may interact weakly with Csd7. Additionally, CcmA localizes to the major helical axis of *H. pylori* cells where rates of PG synthesis are increased (Blair *et al*., 2018; Taylor *et al*., 2020; Yang *et al*., 2019). In this study, we identified several new mechanisms that clarify how CcmA regulates the helical cell shape of *H. pylori* by modulating interactions with Csd5 and Csd7.

First, we identified that the N-terminal region of CcmA and the bactofilin domain are required for helical cell shape, but that the C-terminal region was not required. Next, in co-immunoprecipitation experiments, we found that the bactofilin domain of CcmA interacts with Csd5 and Csd7 and the N-terminal region inhibits the bactofilin domain from interacting with Csd7. When the N-terminal region of CcmA is removed, the bactofilin domain of CcmA binds to Csd7 and Csd1 levels are depleted, providing the first potential mechanism by which CcmA regulates Csd1.

Next, we identified that when other cell-shape-determining genes are deleted, CcmA can no longer localize to the major helical axis of the cell (Gaussian curvature value of 5 ±1 µm^-2^ (Taylor *et al*., 2020)) suggesting that CcmA is directed to its preferred location by a binding partner and is not capable of localizing to or sensing cell envelope curvature on its own. Loss of Csd5 causes substantially less CcmA to localize to the cell envelope, instead CcmA accumulates in the center of cells. The CcmA signal that does co-localize with the cell envelope in Δ*csd5* and Δ*csd5/7* cells is enriched near zero Gaussian curvature, and is depleted at all other Gaussian curvature values. When *csd6* or *csd7* are deleted, CcmA localizes to regions of negative Gaussian curvature. These data suggest that Csd5 recruits CcmA to the cell envelope. If helical cell shape is disrupted (due to loss of another cell shape protein) while Csd5 is still present, Csd5 directs CcmA to regions of negative Gaussian curvature. Additionally, the finding that CcmA localization phenotypes are the same in a Δ*csd5* mutant and a Δ*csd5/7* double mutant suggest that Csd5, not Csd7, controls CcmA localization. These results also support the hypothesis that there are two helical-cell-shape-generating protein complexes, one containing Csd5 and one containing Csd7, and that CcmA may regulate interactions between them (Figure 8).

**Figure 8.**
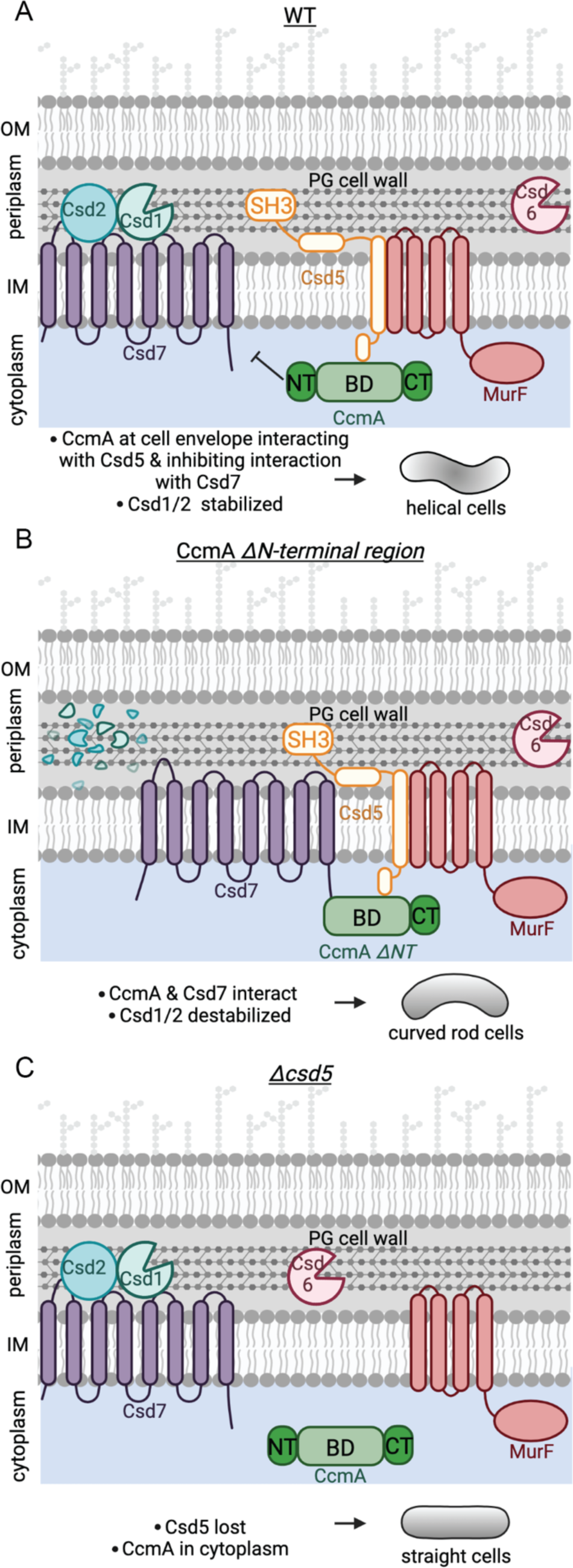
Schematic depicting CcmA’s role in the helical cell shape complexes. There are two protein complexes, one containing Csd5, MurF, and CcmA, and another containing Csd7, Csd1, and Csd2. (A) In WT cells, Csd5 recruits WT CcmA to the cell envelope via CcmA’s bactofilin domain. The N-terminal region of CcmA inhibits interaction with Csd7, allowing Csd1 to function and excluding Csd1 from the CcmA-Csd5-MurF complex. (B) In cells expressing CcmA Δ*NT,* Csd5 recruits Δ*NT* CcmA to the cell envelope via CcmA’s bactofilin domain. The bactofilin domain of CcmA binds to Csd7, inhibiting Csd7 from stabilizing Csd1. (C) When Csd5 is absent, CcmA cannot localize to the cell envelope, MurF is not directed to a particular location. OM = outer membrane, IM =inner membrane. This figure was created using Biorender.com.

Using what we have learned in this study, we can build a new model of the protein complexes that pattern the helical shape of *H. pylori* cells. In one protein complex, CcmA’s bactofilin domain interacts with the cytoplasmic N-terminal region of the transmembrane protein Csd5 directing CcmA to the cell envelope. Csd5 brings MurF and PG precursors to the same region of the cell envelope for new PG to be inserted into the cell wall (Figure 8). CcmA’s N-terminal region inhibits interaction of CcmA’s bactofilin domain with allows Csd7 to stabilize Csd1. When the N-terminus of CcmA is removed, interactions between Csd7 and CcmA are stabilized and Csd1 is no longer detected in the Csd7 complex. Prior work had shown that Csd1 protein cannot be detected in a Δ*csd7* mutant, suggesting Csd7 binding to Csd1 stabilizes this PG hydrolase. Steady state levels of Csd1 are depleted in the *ccmA !1NT* mutant suggesting that Csd7 cannot simultaneously interact with Csd1 and CcmA. Thus, CcmA regulates Csd1 activity (cleaving tetra-pentapeptide crosslinks) spatially or temporally in relation to Csd5 and MurF to organize PG modification and insertion. Our work raises the question of how Csd5 localizes to the major helical axis, Csd5 may sense curvature. Alternatively, the C-terminal SH3 domain of Csd5 can bind to PG directly (Blair *et al*., 2018) and may sense a particular curvature or structure of the cell wall.

In addition to understanding how the helical cell shape of *H. pylori* is controlled, this study provides information about how CcmA and potentially other bactofilins function. Modeling combined with circular dichroism data show that the bactofilin domain is a β-helix and the two short terminal regions are largely unstructured. Additionally, we identified that two previously studied CcmA point mutants that cause loss of function are unfolded (Figure 3). The I55 and L110 residues are homologous to residues mutated by Vasa and colleagues in BacA (V75 and F130 in BacA) and are predicted to reside in the interior of the hydrophobic core of the proteins and mutating these residues destabilizes the proteins likely through the hydrophobic effect (Taylor *et al*., 2020; Vasa *et al*., 2015). We also identified that the bactofilin domain is capable of polymerizing on its own, the N-terminal region promotes lateral interactions between filaments, and the N- and C-terminal regions are both necessary for lattice formation. These data correlate with what has been previously reported in the literature; in the bactofilin Ttbac from *T. thermophilus* loss of the N-terminal region (residues 1-10 in Ttbac) generated smaller, more ordered bundles of filaments (Deng *et al*., 2019), and it has been previously reported that purified Δ*CT* CcmA from *H. pylori* does not form lattice structures (Holtrup *et al*., 2019).

In this study, we identified several putative functional motifs within CcmA. The N-terminal region (amino acids 1-18) is important for patterning helical cell shape and regulates interactions with Csd7, suggesting that the N-terminal region is a functional motif. Our data also suggest that amino acids 13-17 in the N-terminal region of CcmA promote lateral interactions between filaments *in vitro* and may be another functional motif. However, in CcmA, the putative membrane binding motif (amino acids 3-12) suggested by Deng and colleagues (Deng *et al*., 2019), did not show any clear function. Additionally, the C-terminal region of CcmA is not required for any of the cellular phenotypes assayed (cell shape, Csd5 or Csd7 interactions, Csd1 stabilization).

Bactofilins are commonly found as part of complexes in other bacteria and help spatially organize cells to regulate where certain functions occur (Sichel & Salama, 2020; Surovtsev & Jacobs-Wagner, 2018). Our findings regarding how CcmA functions within a complex in *H. pylori* provides insight into how bactofilins operate within complexes in other bacterial species. Mechanisms similar to how CcmA functions in *H. pylori* have been reported in other systems; in *A. biprostecum* a PG hydrolase, SpmX, directs the bactofilin BacA to the site of stalk synthesis, then BacA organizes the rest of the complex (Caccamo *et al*., 2020) to control stalk morphogenesis. Here, we identified that CcmA is not capable of localizing to the cell envelope without Csd5. While Csd5 is unique to the *Helicobacter* genus, transmembrane proteins in other species may also be essential for localization of bactofilins. Interestingly, the genomic arrangement of *csd1* (an M23 metallopeptidase) and *CcmA* overlapping is conserved in several other curved and helical species of Campylobacterota and Pseudomonadota (Billini *et al*., 2019; Caccamo *et al*., 2020; Sycuro *et al*., 2010), and has been reported in a spirochete (Jackson *et al*., 2018), suggesting that bactofilins may regulate M23 metallopeptidases to promote PG remodeling at specific regions of the cell envelope to control cell shape and morphogenesis of other morphological features. Our work is the first to suggest a mechanism by which a bactofilin can regulate a PG hydrolase indirectly through a transmembrane membrane protein and protein stability.

## Materials and Methods

### Bacterial strains and growth

Strains used in this work, as well as primers and plasmids used in strain construction are available in the appendix. *H. pylori* was grown in Brucella Broth (BD) supplemented with 10% heat-inactivated fetal bovine serum (Cytiva Hyclone) without antimicrobials or on horse blood agar plates with microbials as described (Sycuro *et al*., 2010). For resistance marker selection, horse blood agar plates were supplemented with 15 µg/ml chloramphenicol, 25 µg/ml kanamycin, or 60 mg/ml sucrose. *H. pylori* strains were grown at 37 °C under micro-aerobic conditions in a tri-gas incubator (10% CO_2_, 10% O_2_, and 80% N_2_). For plasmid selection and maintenance in *E. coli,* cultures were grown in Lysogeny broth (LB) or on LB-agar at 37 °C supplemented with 100 µg/mL ampicillin or 25 µg/ml Kanamycin.

### Protein structure prediction

Swiss-Model was used to identify the boundaries between the bactofilin domain and the terminal regions of CcmA. The primary amino acid sequence of CcmA from the *H. pylori* G27 genome (GenBank: CP001173.1) was submitted to Swiss-Model (https://swissmodel.expasy.org/ (Waterhouse *et al*., 2018)) where the structure of BacA from *C. crescentus* (Vasa *et al*., 2015) was used to model CcmA. RoseTTAFold was used to predict the structure of CcmA; the same primary amino acid sequence of CcmA was submitted to the Robetta protein prediction service (https://robetta.bakerlab.org/ (Baek *et al*., 2021)). Models generated in RoseTTAFold were modified in Pymol (The PyMOL Molecular Graphics System, Version 2.4.2 Schrödinger, LLC.).

### Western blotting

Whole cell extracts of *H. pylori* were prepared by harvesting 1.0 optical density at 600 nm (OD600) of log phase liquid culture (0.3-0.7 OD600) by centrifugation for 2 minutes. Then, bacteria were resuspended to a concentration of 10 OD600 per ml with 2x protein sample buffer (0.1 M Tris-HCl, 4% SDS, 0.2% bromophenol blue, 20% glycerol) with or without 5% β-mercaptoethanol (BME). Samples were loaded onto Mini-PROTEAN TGX 4-15% polyacrylamide gels (Bio-Rad) and transferred onto polyvinylidene difluoride (PVDF) membranes using the Trans-Blot TurboTransfer System (Bio-Rad) system. Membranes were either stained with 0.1% Ponceu S solution in 5% acetic acid (Sigma) first, or immediately blocked for 2 hours at room temperature in 5% non-fat milk in tris-buffered saline containing 0.05% tween-20 (TBS-T) then incubated either overnight at 4 °C or at room temperature for 1 hour with primary antibodies, 1:10,000 for ɑ-CcmA (Blair et al., 2018), 1:5,000 for ɑ-FLAG (M2, Sigma), 1:10,000 for ɑ-Csd1 (Yang *et al*., 2019), 1:20,000 for ɑ-cag3 (Pinto-Santini & Salama, 2009), or 1:10,000 for ɑ-Csd1 (Promega). Then washed 5 times with TBS-T over a 30-minute period, incubated with secondary antibodies for 1 hour at room temperature at 1:20,000 with appropriate horseradish peroxidase-conjugated ɑ-immunoglobulin antibodies (ɑ-rabbit for CcmA Csd1, or Cag3; ɑ-mouse for FLAG, Santa Cruz Biotechnology), then washed 5 times with TBS-T over a 30-minute period. Proteins were detected using a chemiluminescent substrate according to manufacturer’s instructions (Pierce ECL substrate for standard Western blotting or Immobilon Western HRP substrate (Millipore) for Western blotting following co-immunoprecipitation experiments) and imaged directly with the Bio-Rad gel documentation system.

### Western blot densitometry

Western blot densitometry was performed using Image Lab software version 6.0.1 (Bio-Rad). The lanes on each Western blot were defined, then the adjusted total lane volume was calculated by the software. To calculate relative Csd1 levels, the signal detected by the ɑ-Csd1 antibody for each sample was divided by the signal detected by the ɑ-Cag3 antibody from the same sample on the same membrane. After calculating relative Csd1 levels, they were normalized by dividing the signal by the relative Csd1 level from the SSH51A strain (*rdxA::csd1*).

### Phase contrast microscopy and quantitative morphology analysis

Phase contrast microscopy and image analysis was performed as previously described (Sycuro *et al*., 2010). Briefly, cells were grown in liquid culture to mid-log phase (0.3-0.7 OD_600_), centrifuged, and resuspended in 4% paraformaldehyde (PFA) in phosphate-buffered saline (PBS) and mounted on glass slides. Cells were imaged with a Nikon TE 200 microscope equipped with a 100X oil immersion lens objective and Photometrics CoolSNAP HQ charge-coupled-device camera. Images were thresholded with FIJI (Schindelin *et al*, 2012) and CellTool software (Pincus & Theriot, 2007) was used to measure both side curvature and central axis length.

### Protein purification

Purification of 6-his CcmA was performed exactly as described previously (Taylor *et al*., 2020). *E. coli* strains used to recombinantly express CcmA were: pKB62A in BL21 DE3, pSS15A in BL21 DE3, pSS16A in BL21 DE3, pSS17A in BL21 DE3, pSS18B in BL21 DE3, pSS19A in BL21 DE3, pKB69H in BL21 DE3, and pKB72D in BL21 DE3 and are described in detail in the appendix.

### Circular dichroism

After purification of 6-his CcmA, samples were dialyzed against 10 mM CAPS pH 11 at a volume of 1:10,000 for 24 hours, then the buffer was changed, and dialysis continued for another 24 hours. After dialysis was complete, samples were normalized to 0.5 mg/ml. CD spectra were collected from 180-260 nm while holding the temperature constant with a Jasco J-815 Circular Dichroism Spectropolarimeter.

### Negative stain and Transmission Electron Microscopy (TEM)

After purification of 6-his CcmA samples were dialyzed against 25 mM Tris pH 8 for 24 hours. Samples were then normalized to 1 mg/ml and placed on carbon-coated grids after glow-discharging. Samples were fixed in ½ X Karnovsky’s fixative and washed with 0.1 M Cacodylate buffer and water, then stained with Nano-W (Nanoprobes) for 60 seconds, wicked, then stained for another 60 seconds. Images were acquired on a kV ThermoFisher Talos 120C LaB6 microscope equipped with a Ceta Camera using Leginon software (Suloway *et al*, 2005). The width of bundles and filaments were measured using FIJI (Schindelin *et al*., 2012).

### Co-immunoprecipitation experiments

Co-immunoprecipitation experiments were performed as previously described in Blair *et al.,* 2018 and Yang *et al.,* 2019 with slight modifications. *H. pylori* expressing FLAG tags fused to either Csd5 or Csd7 were grown to mid-log phase in liquid culture and 15 OD_600_ were harvested by centrifugation at 10,000 RPM for 10 minutes at 4 °C. Cells pellets were resuspended in 1.8 ml of cold lysis buffer (20 mM tris pH 8, 150 mM NaCl, 2% glycerol, 1% Triton X-100 and EDTA-free protease inhibitor (Pierce)). The cells were sonicated at 10% power for 10 second intervals with a microtip (Sonic Dismembrator Model 500, Branson) until visibly cleared, then centrifuged at 20,000 RCF for 30 minutes at 4 °C. The soluble fraction was then incubated with 40 µl beads (ɑ-FLAG M2 agarose beads (Sigma)) or ɑ-VSV-G agarose beads (Sigma) equilibrated in wash buffer (20 mM tris pH 8, 150 mM NaCl, 2% glycerol, 0.1% Triton X-100) for 90 minutes at 4 °C. The beads were then washed four times in 10 ml cold wash buffer, samples were centrifuged at 2000 RPM for 2 minutes between washes. Beads were then boiled in 45 µl 2x sample buffer without BME for 10 minutes and subjected to Western blotting.

### Labeling with fluorescent WGA and HaloTag ligands

*H. pylori* expressing CcmA-HaloTag were grown to mid-log phase and fixed with 4% PFA for 45 minutes at room temperature. After fixing, bacteria were permeabilized for 1 hour at room temperature with PBS + 0.05% Triton X-100. Bacteria were then resuspended in PBS and centrifuged onto #1.5 coverslips. Next, coverslips were incubated on a droplet of PBS containing 30 µg/ml af488-conjucated WGA (Invitrogen) and 4 µM JF-549 (Promega) for 30 minutes at room temperature. Coverslips were washed four times (10 minutes each) by placing on fresh droplets of PBS + 0.1% Tween-20, then mounted on glass slides with ProLong Diamond Antifade Mountant (Invitrogen).

### 3D Structured Illumination Microscopy

3D structured illumination microscopy (SIM) was performed on a DeltaVIsion OMX SR equipped with PCO scientific CMOS cameras, 488 nm and 568 nm lasers, and an Olympus 60X/1.42 U PLAN APO oil objective with oil matched for the sample refractive index. 512 x 512 pixel Z-stack images were acquired with 125 nm spacing and 3 µm total thickness. Figures were generated by changing channel colors, adjusting brightness and contrast in FIJI, brightness and contrast settings were maintained across the entire image (Schindelin *et al*., 2012). Figures were assembled in Adobe Illustrator.

### 3D reconstructions and curvature enrichment analysis

3D reconstructions were generated, and curvature CcmA enrichment analysis was preformed exactly as described in Taylor *et al.,* 2020 using existing software (Bratton *et al*., 2019; Bratton *et al*, 2018). Briefly, 3D SIM Z-stack images were used to generate a 3D triangular meshwork surface with roughly 30 nm precision from the WGA channel. After generating the 3D triangular meshwork, the Gaussian curvature was calculated at every location on the surface of the cell and we performed curvature enrichment analysis to identify which GC values CcmA signal was associated with where uniform signal=1. Because the absolute amount of CcmA signal can differ between cells and because the illumination throughout the field is non-uniform, we set the average CcmA signal on each cell as one. Then we measured each cell’s curvature dependent CcmA signal intensity relative to one, normalized by the amount of the curvature present on that individual cell surface.

### Analysis of intensity of CcmA-HaloTag signal at cell surface and within cells

A custom Matlab script was used to compute the relative intensity of CcmA-HaloTag signal at the cell surface and at surfaces generated by shrinking the surface of interest toward the centerline of the cell (negative offsets) or increasing the surface away from the centerline (positive offsets). For each of these shells, or surface offsets, we calculated the average WGA signal and the average CcmA-HaloTag signal. To account for errors in spatial alignment between the two color channels, before calculating the intensity at each shell, we performed a rigid body alignment procedure to maximize the intensity of the signal contained within the nominal reconstructed surface (0 nm offset). Consistent with the fact that we utilized the WGA channel for shape reconstructions, this alignment algorithm returned offsets of less than one pixel for the WGA channel. The CcmA-HaloTag channel showed more variability in the offsets, sometimes requiring an offset of almost 200 nm along the focal direction, consistent with small, wavelength dependent variations in focal position. After calculating the average intensity at each subshell, or resized surface, we normalized the highest concentration to 1 before averaging across all the cells in our sample.

## Supporting information

Appendix containing Supplemental Tables and Methods

## Acknowledgements

This research was supported by the US National Institute of Heath (NIH) grants F31 AI152331 (SRS), T32 GM95421 (SRS), R01 AI136946 (NRS). As well as the GO-MAP Graduate Opportunity Program Research Assistantship award (GOP Award) (SRS) and the VUMC Discovery Scholars in Health and Medicine Program (BPB). This research was supported by the Genomics and Cellular Imaging and Electron Microscopy shared resources of the Fred Hutch/University of Washington Cancer Consortium (P30 CA015704). The opinions, findings, and conclusions or recommendations expressed in this material contents are solely the responsibility of the authors and do not necessarily represent the official views of the NIH.

We would like to thank Joseph Stembel (University of Washington) for construction of strains JS02 and JS09, Jennifer Taylor (University of Washington) for helpful advice and training, Ethan Garner (Harvard) for helpful discussion, James McDermott for thoughtful discussion and review of this manuscript, David Baker and Justin Decarreau (University of Washington), and the support of the Audacious Project at the Institute for Protein Design for use of the DeltaVision OMX microscope, Roland Strong and Barry Stoddard (Fred Hutchinson Cancer Center) for use of the Circular Dichroism Spectropolarimeter, and Caleigh Azumaya (Fred Hutchinson Cancer Center) for assistance with the Talos microscope.

## Data Availability

The datasets and computer code produced in this study will be available soon at the following locations:

- Matlab scripts and documentation: https://github.com/PrincetonUniversity/shae-cellshape-public.
- Imaging datasets: https://www.ebi.ac.uk/bioimage-archive/

## Author contributions

Conception and design of the study (SRS, NRS), acquisition of data (SRS), analysis and interpretation of data (SRS, NRS, BPB), writing the manuscript (SRS, NRS).

## Supplemental Figures

**Supplemental Figure 1.**
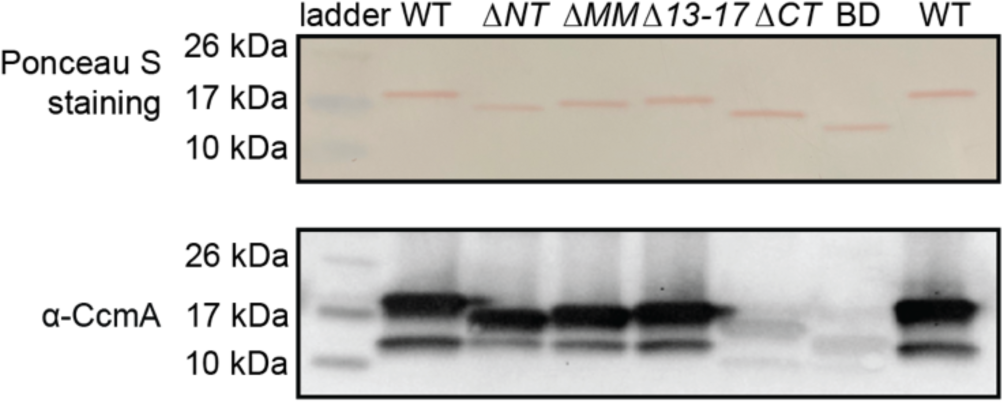
Robust CcmA detection with ɑ-CcmA polyclonal antibody requires the C-terminus. 0.1 µg of purified protein was loaded into each lane. Top row: Ponceau S staining of purified 6-his-CcmA. Bottom Row: Western blot probed with polyclonal ɑ-CcmA antibody, both images are of the same membrane.

**Supplemental Figure 2.**
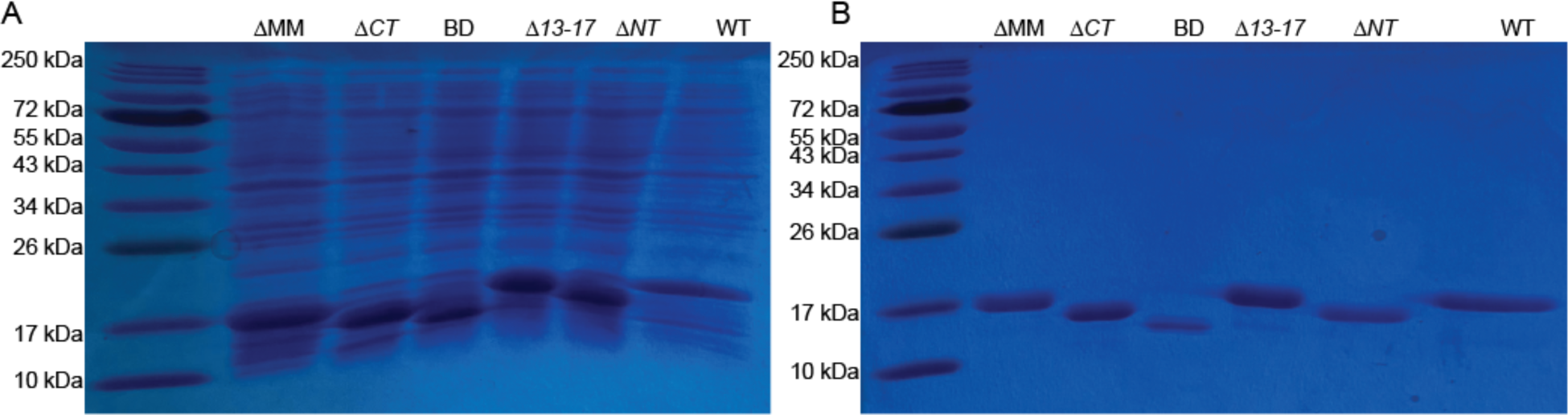
Purification of WT and mutant 6-his CcmA. (A) Coomassie-stained gel of induced, whole cell lysates of *E. coli*. (B) Coomassie-stained gel of the first elution after purification of 6-his CcmA with cobalt affinity resin.

**Supplemental Figure 3.**
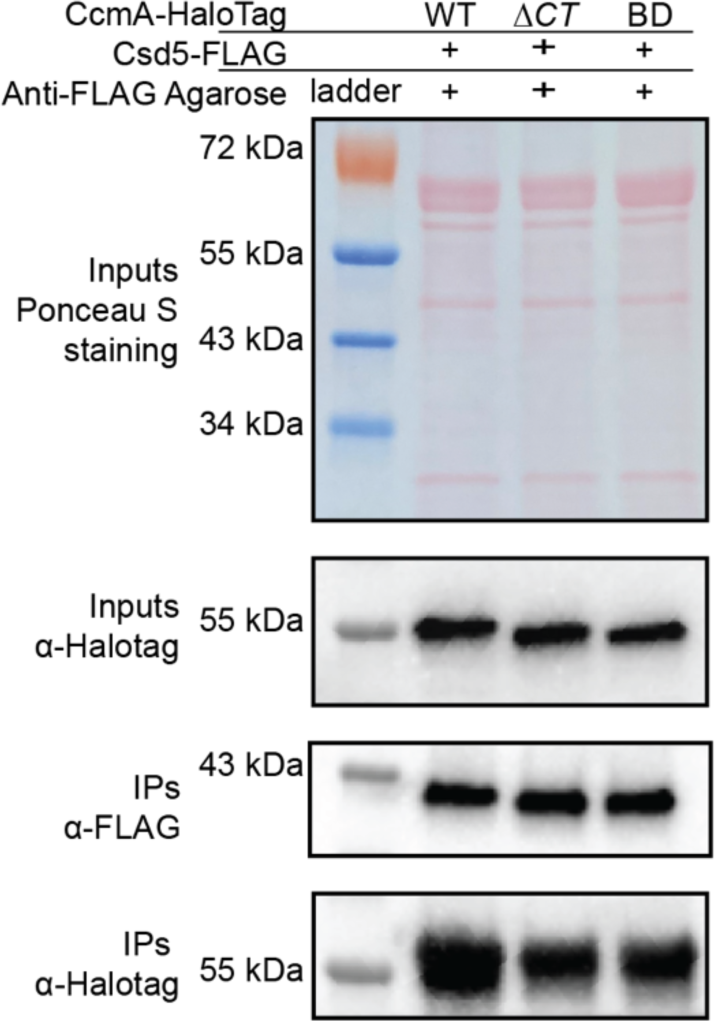
The C-terminus of CcmA-HaloTag is not required for interactions with Csd5. ɑ-FLAG immunoprecipitation of strains expressing Csd5-FLAG and WT CcmA-HaloTag, CcmA Δ*CT*-HaloTag, or CcmA BD only -HaloTag. Top Row: Ponceau S staining of input fractions. Second row: Western blot of inputs probed with ɑ-HaloTag monoclonal antibody to detect CcmA-HaloTag, both top and second row are images of the same membrane. Third row: Western blot of IP fractions probed with ɑ-FLAG monoclonal antibody to detect Csd5-FLAG. Bottom row: Western blot of IP fractions probed with ɑ-HaloTag monoclonal antibody to detect CcmA-HaloTag. Strains used: SSH87A, SSH89A, and SSH97A

**Supplemental Figure 4.**
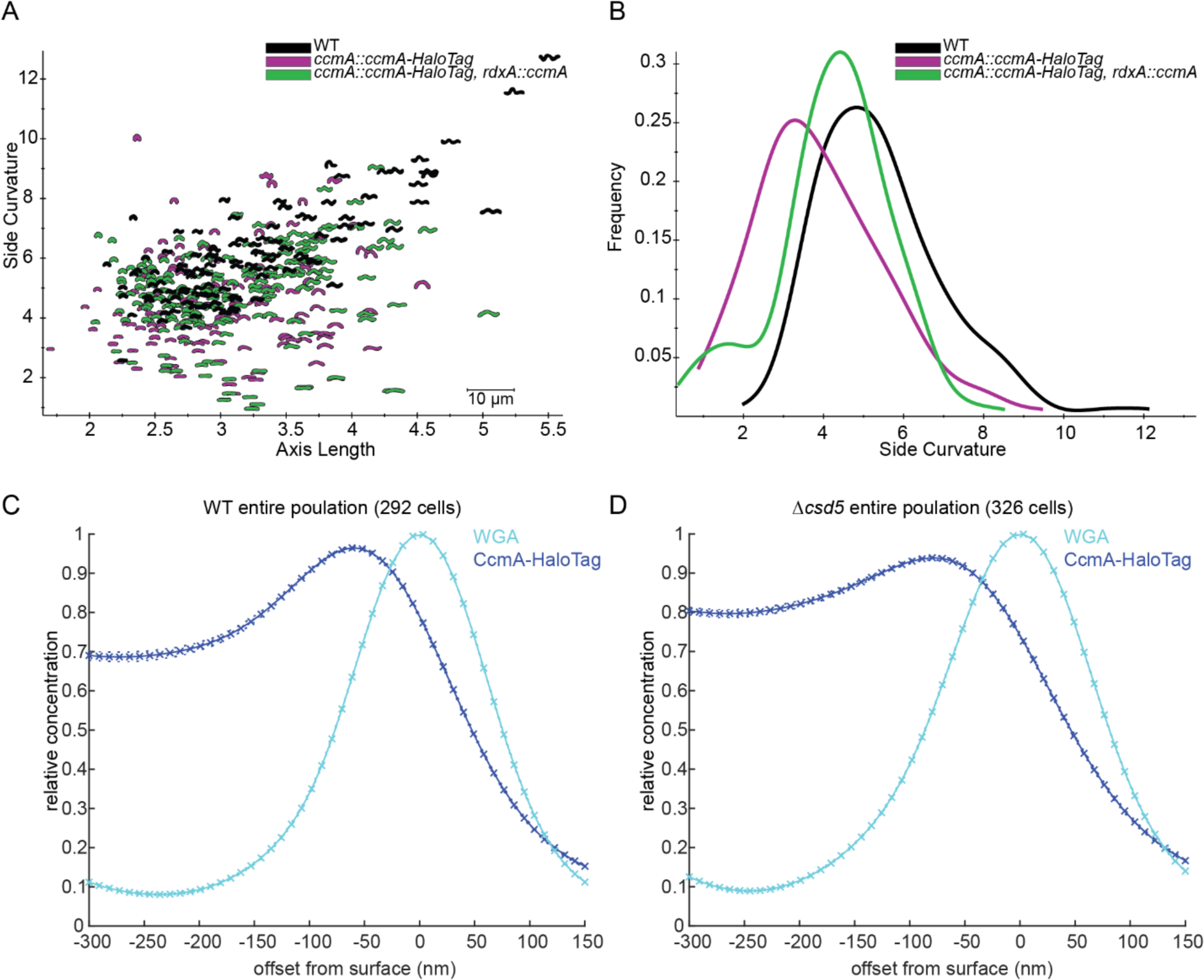
Cell shape of CcmA-HaloTag strains and analysis of CcmA-HaloTag surface signal in populations. (A) Scatterplot of outlines of *H. pylori* cells derived from phase contrast images. (B) Histogram displaying side curvature of each population. WT (LSH100) n=116 cells, *ccmA::ccmA-Halotag* (SSH38A) n=133 cells, *ccmA::ccmA-HaloTag, rdxA::ccmA* (SSH39B) n=190 cells. (C-D) Plots displaying the relative concentration of HaloTag-CcmA and WGA signal at the cell surface (X=0 nm) and at computationally generated cell surfaces that are inside (X<0 nm) and outside (X>0 nm) of the cell surface. Crosses indicate SEM. Populations of WT (C) and Δ*csd5* (D) cells from three independent experiments were pooled.

