## Appendix containing Supplemental Tables and Methods for "Distinct regions of *H. pylori’s* bactofilin CcmA regulate protein-protein interactions to control helical cell shape"

Table S1. *H. pylori* strains used in this work
Table S2. *E. coli* strains used in this work
Table S3. Plasmids used in this work
Table S4. Primers used in this work

**Supplemental experimental procedures**

Genetic manipulation of *H. pylori* strains and construction of *E. coli* vectors

**Supplemental references**

**BioRender Confirmation of Publication and Licensing Rights**

**Table S1. *H. pylori* strains used in this work**

| **Strain** | **Relevant genotype or Description** | **Reference or source** |
|---|---|---|
| DCY26 | *csd7::catsacB* | (Yang *et al*, 2019) |
| DCY28 | *csd7::cat* | (Yang *et al.*, 2019) |
| DCY71 | *csd7-3X-FLAG* | (Yang *et al*, 2019) |
| JS02 | *ccmA::ccmA-40 aa linker-HaloTag, rdxA::ccmA, csd5::cat2kan* | This work |
| JS09 | *ccmA::ccmA-40 aa linker-HaloTag, rdxA::ccmA, csd5::cat2kan, csd7::cat* | This work |
| JTH2 | *csd5-2X-FLAG cat* | (Blair *et al*, 2018) |
| JTH3 | *ccmA-2X-FLAG cat2kan* | (Blair *et al.*, 2018) |
| KBH157 | *csd5-2X-FLAG, murF-3X-VSV-G::cat* | (Blair *et al.*, 2018) |
| LSH100 | Wild-type *H. pylori*, NSH57 with *fliM* repaired | (Sycuro *et al*, 2010) |
| LSH108 | *rdxA::kansacB* | (Sycuro *et al.*, 2010) |
| LSH113 | *csd1::cat* | (Sycuro *et al.*, 2010) |
| LSH117 | *ccmA::catsacB* | (Sycuro *et al.*, 2010) |
| LSH121 | *csd1::cat, rdxA::csd1* | (Sycuro *et al.*, 2010) |
| LSH123 | *csd5::cat* | (Sycuro *et al*, 2012) |
| LSH148 | *ccmA::catsacB, rdxA::ccmA* | (Sycuro *et al.*, 2010) |
| SSH31C | *ccmA::ccmA-12 aa linker-HaloTag* | This work |
| SSH33 | *ccmA::ccmA-40 aa linker-mNeonGreen* | This work |
| SSH38A | *ccmA::ccmA-40 aa linker-HaloTag* | This work |
| SSH39B | *ccmA::ccmA-40 aa linker-HaloTag, rdxA::ccmA* | This work |
| SSH41A | *ccmA::catsacB*, *csd7::csd7-3X-FLAG* | This work |
| SSH49A | *ccmA::ccmA-40 aa linker-HaloTag, rdxA::ccmA, csd5::cat* | This work |
| SSH50A | *ccmA::ccmA-40 aa linker-HaloTag, rdxA::ccmA, csd7::catsacB* | This work |
| SSH51A | *rdxA::csd1* | This work |
| SSH53A | *ccmA::catsacB, rdxA::csd1* | This work |
| SSH54A | *ccmA::ccmA ∆MM, rdxA::csd1* | This work |
| SSH55A | *ccmA::ccmA ∆NT, rdxA::csd1* | This work |
| SSH56B | *ccmA::ccmA ∆CT, rdxA::csd1* | This work |
| SSH57A | *ccmA::ccmA BD, rdxA::csd1* | This work |
| SSH58A | *ccmA::ccmA ∆MM, rdxA::csd1, csd5::csd5-2X-FLAG-cat* | This work |
| SSH59A | *ccmA::ccmA ∆NT, rdxA::csd1, csd5::csd5-2X-FLAG-cat* | This work |
| SSH60A | *ccmA::ccmA ∆CT, rdxA::csd1, csd5::csd5-2X-FLAG-cat* | This work |
| SSH65A | *ccmA::ccmA BD, rdxA::csd1, csd5::csd5-2X-FLAG-cat* | This work |
| SSH67A | *ccmA::ccmA ∆13-17, rdxA::csd1* | This work |

| | | |
|---|---|---|
| SSH68A | *ccmA::ccmA ∆13-17, rdxA::csd1, csd5::csd5-2X-FLAG-cat* | This work |
| SSH70A | *ccmA::ccmA-40 aa linker-HaloTag, rdxA::ccmA, csd6::cat* | This work |
| SSH78B | *ccmA::ccmA ∆MM, rdxA::csd1, csd7::csd7-3X-FLAG* | This work |
| SSH79A | *ccmA::ccmA ∆NT, rdxA::csd1, csd7::csd7-3X-FLAG* | This work |
| SSH80A | *ccmA::ccmA ∆CT, rdxA::csd1, csd7::csd7-3X-FLAG* | This work |
| SSH81B | *ccmA::ccmA BD, rdxA::csd1, csd7::csd7-3X-FLAG* | This work |
| SSH82 | *ccmA::ccmA ∆13-17, rdxA::csd1, csd7::csd7-3X-FLAG* | This work |
| SSH87A | *ccmA::ccmA-40 aa linker-HaloTag, csd5::csd5-2X-FLAG-cat* | This work |
| SSH89A | *ccmA::ccmA ∆CT-40 aa linker-HaloTag, rdxA::csd1,csd5::csd5-2X-FLAG-cat* | This work |
| SSH97A | *ccmA::ccmA BD-40 aa linker-HaloTag, rdxA::csd1, csd5::csd5-2X-FLAG-cat* | This work |
| TSH17 | *csd6::cat* | (Sycuro *et al*, 2013) |

**Table S2*. E. coli* strains used in this work**

| **Strain** | **Relevant genotype or Description** | **Reference or source** |
|---|---|---|
| BL21 (DE3) | Protein expression *E. coli* strain | New England Biolabs |
| Stellar | Cloning strain of *E. coli* | Takara |

**Table S3. Plasmids used in this work**

| **Plasmid** | **Genotype or Description** | **Marker** | **Reference or Source** |
|---|---|---|---|
| pET15b | Modified pET15 vector (6-his expression vector) | Ampicillin | Barry Stoddard Lab, Fred Hutchinson Cancer Center |
| pKB62A | *6-his ccmA* in pET15b | Ampicillin | (Taylor *et al*, 2020) |
| pLC292 | pRdxA | Ampicillin | (Terry *et al*, 2005) |
| pLKS31 | *csd1* in pLC292 | Ampicillin | (Sycuro *et al.*, 2010) |
| pSS15A | *6-his ccmA ∆MM* in pET15b | Ampicillin | This work |
| pSS16A | *6-his ccmA ∆CT* in pET15b | Ampicillin | This work |
| pSS17A | *6-his* ccm*A BD* in pET15b | Ampicillin | This work |

| | | | |
|---|---|---|---|
| pSS18B | *6-his ccmA ∆13-17* in pET15b | Ampicillin | This work |
| pSS19A | *6-his ccmA ∆NT* in pET15b | Ampicillin | This work |
| pCR Blunt II-TOPO vector | TOPO cloning vector | Kanamycin | Invitrogen |
| pKB69H | *6-his ccmA I55A* in pET15b | Ampicillin | (Taylor *et al.*, 2020) |
| pKB72D | *6-his ccmA L110S* in pET15b | Ampicillin | (Taylor *et al.*, 2020) |
| pFC30K | His6HaloTag T7 Flexi Vector | Kanamycin | Promega |
| pSS6-4 | *ccmA-12 aa linker-HaloTag* in pCR Blunt II-TOPO | Kanamycin | This work |
| pSS8E | *ccmA-40 aa linker-mNeonGreen* in pCR Blunt II-TOPO | Kanamycin | This work |
| pSS10-1 | *ccmA-40 aa linker-HaloTag* in pCR Blunt II-TOPO | Kanamycin | This work |

**Table S4. Primers used in this work**

| **Name** | **Sequence 5' to 3'** |
|---|---|
| **Construction of *H. pylori* strains** | |
| Csd1 F | GAGTCGTTACATTAATGTGCATATCT |
| CcmA SDM dn R | GCTCATTTGAGTGGTGGGAT |
| remove aa 2-17 F | CAATAAAGAAAGGAGCATCAGATGGCGACTATCATCGCTC |
| remove aa 2-17 R | GAGCGATGATAGTCGCCATCTGATGCTCCTTTCTTTATTG |
| remove aa 3-12 F | AATAAAGAAAGGAGCATCAGATGGCAGCAAAAACAGG |
| remove aa 3-12 R | CCTGTTTTTGCTGCCATCTGATGCTCCTTTCTTTATT |
| remove aa 119-136 F | GGGAAACTCGCCCTAAGAATTAGGGAATGATCCAATCTAG |
| remove aa 119-136 R | CTAGATTGGATCATTCCCTAATTCTTAGGGCGAGTTTCCC |
| remove aa 13-17 2 F' | ATAACAATAATAAATCGGCTAATGCGACTATCATCGCTCA |
| remove aa 13-17 2 R' | TGAGCGATGATAGTCGCATTAGCCGATTTATTATTGTTAT |
| CcmA_40aa_linker_upstream_R | ACCTTGTCCGCTACCCTCAAGTTTATTTTCAATTTTCTTTTC |
| CcmA_mNeonGreen_dnstrm_F | GATGGATGAATTATATAAATAATAGGGAATGATCCAATCTAGTCT |

| | |
|---|---|
| CcmA_12_aa_link_Halo_R | AGTACCGATTTCACTACCACCACCACTACCACCACCACTACCACCACCTTTATTTTCAAT |
| CcmA_Halo_dnstrm_R | AGACTAGATTGGATCATTCCCTAACCGGAAATCTCCAGA |
| CcmA_12_aa_link_Halo_F | ATTGAAAATAAAGGTGGTGGTAGTGGTGGTGGTAGTGGTGGTGGTAGTGAAATCGGTACT |
| 40 aa linker halotag R | TGGAAAGCCAGTACCGATTTCGCCTTGACCTGGGCCAGATCC |
| 40 aa linker halotag F | GGATCTGGCCCAGGTCAAGGCGAAATCGGTACTGGCTTTCCA |
| C1_828 | GATATAGATTGAAAAGTGGAT |
| C1_829 | TTATCAGTGCGACAAACTGGG |
| **Cloning for protein purification** | |
| CcmA_pET15b_BamHI | GTTAGCAGCCGGATCCCTATTTATTTTCAATTTTCTTTTCTTGCTCATTGA |
| pET15b_ccmA_delta_aa2to17_NdeI | CGCGCGGCAGCCATATGATGGCGACTATCATCGCTCAAG |
| CcmA_pET15b_BamHI | GTTAGCAGCCGGATCCCTATTTATTTTCAATTTTCTTTTCTTGCTCATTGA |
| pET15b_ccmA_delta_aa3to12_NdeI | CGCGCGGCAGCCATATGATGGCAGCAAAAACAGGACCAG |
| CcmA_pET15b_NdeI | CGCGCGGCAGCCATATGATGGCAATCTTTGATAACAATAATAAATCGGCT |
| pET15b_ccmA_delta_aa119to136_BamHI | GTTAGCAGCCGGATCCCTAATTCTTAGGGCGAGTTTCC |
| pET15b_ccmA_delta_aa13to17_NdeI | CGCGCGGCAGCCATATGATGGCAATCTTTGATAACAATAATAAATCGGCT |

| **Sequencing primers** | | |
|---|---|---|
| Target | Primer name | |
| *rdxA* | 1318 | GAAGCGGTTACAATCATCACGCCC |
| *rdxA* | 1319 | GCTTGAAAACACCCCTAAAAGAGCG |
| *ccmA* | 1432 | GATTACCATTTGCATGTAGATGGCG |
| *ccmA* | 1433 | CTAGAGATCTTACCATCAAGGCGC |
| *csd5* | Csd5-c-term-seq | AAGCGTGAAGGTTTTAGAAATCCA |
| *csd5* | 1194_R | CCACAAGCTCATCATCTTCCAAAA |
| pET15b | T7 F | CGAAATTAATACGACTCACTATAGG |
| pET15b | T7 R | CCTCAAGACCCGTTTAGAGGCC |

#### Supplemental experimental procedures

##### **<u>Genetic manipulation of *H. pylori* strains and construction of *E. coli* vectors</u>**

All genetic manipulations of *H. pylori* were performed on the chromosome via natural transformation and allelic exchange by homologous recombination of either purified PCR products or plasmids. We used chloramphenicol (*cat*) and kanamycin (*aphA3*) resistance cassettes as markers in some strains (Trieu-Cuot *et al*, 1985; Wang & Taylor, 1990). In others, we used selectable and counter-selectable cassettes *catsacB* (chloramphenicol resistance and sucrose susceptibility) and *kansacB* (kanamycin resistance and sucrose susceptibility) for constructing marker-less strains (Copass *et al*, 1997).

All genomic DNA preparations from *H. pylori* were performed with the Wizard Genomic DNA Purification Kit (Promega). All plasmids were purified with the QIAprep Spin MiniPrep Kit (Qiagen).

<u>Construction of *H. pylori* strains expressing truncated versions of *ccmA*</u>

To construct strains expressing truncated versions of CcmA, we first generated a parent strain expressing an extra copy of *csd1* at *rdxA,* a locus commonly used for complementation (Smeets *et al*, 2000). First, we transformed LSH108 (*rxdA::kansacB*) with plasmid pLKS31 (*csd1* in pRdxA) to replace *kansacB* at *rdxA* with *csd1*, then selected for colonies that grew on horse blood agar (HB) plates (Humbert & Salama, 2008) containing sucrose but could not grow on HB plates containing kanamycin to generate strain SSH51A. Then, we replaced the *ccmA* gene with a *catsacB* cassette so marker-less truncated versions of *ccmA* could be engineered by allelic exchange. To do this we transformed strain SSH51A with genomic DNA from LSH117 (*ccmA::catsacB*) and selected for colonies that grew on HB plates containing chloramphenicol to generate strain SSH53A. We transformed strain SSH53A with purified PCR products generated by PCR-SOEing (Horton, 1995) that had truncated versions of *ccmA* flanked by 800 nucleotides of homology upstream of the *ccmA* and 500 nucleotides of homology downstream of the *ccmA,* and selected for colonies that grew on HB plates containing sucrose, but not on HB plates containing chloramphenicol. After confirming that strains grew on selective plates, strains were confirmed by PCR and Sanger Sequencing to sequence the *rdxA* and *ccmA* loci.

To generate the Δ*NT* strain (SSH55A, lacking amino acids 2-17) we used primers 'Csd1 F' and 'remove aa 2-17 R' to amplify the upstream PCR product and primers 'remove aa 2-17 F' and 'CcmA SDM dn R' to amplify the downstream PCR product from LSH100 (WT) genomic DNA. The resulting products were purified using the QIAquick PCR Purification Kit (Qiagen). Then we stitched them together with PCR SOEing (Horton, 1995), gel extracted the product using the QIAquick Gel Extraction Kit (Qiagen) and used the product directly for natural transformation of SSH53A. To generate the Δ*MM* strain (SSH54A, lacking amino acids 3-12), the same strategy was used, but we used primers 'remove aa 3-12 F' and 'remove aa 3-12 R' instead of 'remove aa 2-17 F' and 'remove aa 2-17 R'. To generate Δ*13-17* strain (SSH67A, lacking amino acids 13-17) the same strategy was used, but with primers 'remove aa 13-17 2 F' and 'remove aa 13-17 2 R'. To generate the Δ*CT* strain (SSH56B, lacking amino acids 119-136), we followed the same strategy but used primers 'remove aa 119-136 F' and 'remove aa 119-136 R'.

To generate the BD strain (SSH57A, only amino acids 18-188), we amplified genomic DNA from strain SSH55A by PCR with primers 'Csd1 F' and 'remove aa 119-136 R' and genomic DNA from SSH56B with primers 'CcmA SDM dn R' and 'remove aa 119-136 F' and purified them using the QIAquick PCR Purification Kit (Qiagen). Then we stitched them together with PCR SOEing (Horton, 1995), gel extracted the product using the QIAquick Gel Extraction Kit (Qiagen) and used the product directly for natural transformation of SSH53A.

<u>Construction of *H. pylori* strains expressing FLAG-tagged versions of Csd5 or Csd7</u>

To construct strains expressing Csd5-2X FLAG*,* strains with truncated versions of *ccmA* (SSH54A, SSH55A, SSH56B, SSH57A, SSH67A) were transformed with genomic DNA from JTH2 (contains a cat cassette downstream of *csd5-2X FLAG* at the native *csd5* locus*)*. Colonies that grew on HB plates containing chloramphenicol were selected and confirmed by PCR. These are strains SSH58A, SSH59A, SSH60A, SSH65A, and SSH68A.

To construct strains expressing Csd7-3X FLAG, DCY71 (Csd7-3X-FLAG) was transformed with genomic DNA from LSH117 to replace *ccmA* with *catsacB* and clones that grew on HB plates supplemented with chloramphenicol were selected to generate strain SSH41A. Then, SSH41A was transformed with genomic DNA from strains SSH54A, SSH55A, SSH56B, SSH57A, or SSH67A to replace *catsacB* with truncated versions of *ccmA*, clones that grew on HB plates supplemented with sucrose but not chloramphenicol were selected. Next, strains were transformed with genomic DNA from LSH108 to place *kansacB* at the *rdxA* locus, clones that grew on HB plates supplemented with kanamycin were selected. Finally, clones were transformed with genomic DNA from SSH51A to replace *kansacB* with *csd1* at *rdxA,* clones that grew on HB plates supplemented with sucrose but not on HB plates supplemented with kanamycin were selected and confirmed by PCR and Sanger sequencing of the *rdxA,* and *ccmA* loci. Confirmed strains are SSH78B, SSH79A, SSH80A, SSH81B, SSH82.

<u>Construction of *H. pylori* strains expressing *CcmA-HaloTag*</u>

We used a flexible 40 amino-acid-long linker to link a HaloTag to the C-terminus of CcmA. To do this, we first constructed a strain where the C-terminus of *ccmA* was fused to *mNeonGreen* with the 40 amino-acid-long linker between the two, then modified it to replace *mNeonGreen* with *HaloTag.* We used a gBlock (IDT) that contained the linker followed by a codon-optimized version of *mNeonGreen* and used PCR SOEing (Horton, 1995) to generate a construct that had *ccmA-linker-mNeonGreen* flanked with 800 nucleotides of homology upstream of the *ccmA* and 500 nucleotides of homology downstream of the *ccmA.* We amplified the upstream portion of the construct from LSH100 (WT) genomic DNA with primers 'Csd1 F' and 'CcmA-40aa_linker_upstream_R' and the downstream portion with primers 'CcmA-mNeonGreen_dnstrm_R' and 'CcmA SDM dn R'. After cleaning up the PCR products with QIAquick PCR Purification Kit (Qiagen), we performed PCR-SOEing (Horton, 1995) to stitch together the two PCR products and the gBlock, the product was gel extracted using QIAquick Gel Extraction Kit (Qiagen), then TOPO cloned using the Zero Blunt TOPO PCR cloning Kit (Invitrogen). The plasmid was then used as a template to transform LSH117 for allelic exchange to replace the *catsacB* cassette at the *ccmA* locus with the construct. Clones that grew on

HB plates containing sucrose but not chloramphenicol were selected and confirmed by PCR and Sanger sequencing to generate strain SSH33.

Next, we constructed a strain where HaloTag was fused to the C-terminus of *ccmA* with a 12-amino-acid-long linker. To do this we generated a construct containing *ccmA-12aa-linker-HaloTag* flanked by 800 nucleotides of homology upstream of the *ccmA* and 500 nucleotides of homology downstream of the *ccmA.* We amplified PCR products from genomic DNA from LSH100 with primers 'Csd1 F' and 'CcmA_12_aa_link_HaloR' and primers 'CcmA_Halo_dnstrm_R' and 'CcmA_SDM_dn_R'. Next, we amplified a PCR product containing HaloTag from plasmid pFC30K (Promega) with primers 'CcmA_12_aa_link_Halo_F' and 'CcmA_Halo_dnstrm_R'. After cleaning up the PCR products with QIAquick PCR Purification Kit (Qiagen), we performed PCR-SOEing (Horton, 1995) to stitch the three PCR products together and TOPO cloned the purified PCR product using the Zero Blunt TOPO PCR cloning Kit (Invitrogen). The plasmid was then used as a template to transform LSH117 and replace the *catsacB* cassette at the *ccmA* locus with the construct to generate strain SSH31C. Clones that grew on HB plates containing sucrose but not chloramphenicol were selected and confirmed by PCR and Sanger sequencing.

Finally, to construct a strain expressing *ccmA-40 aa-linker-HaloTag* at the native *ccmA* locus, we amplified a PCR product with primers 'Csd1 F' and '40 aa linker halotag R' from SSH33 genomic DNA and another PCR product with primers '40 aa linker halotag F' and 'CcmA SDM dn R' from SSH31C genomic DNA. After cleaning up the PCR products with QIAquick PCR Purification Kit (Qiagen), we performed PCR-SOEing (Horton, 1995) to stitch the two products together, then gel extracted using QIAquick Gel Extraction Kit (Qiagen), and TOPO cloned using the Zero Blunt TOPO PCR cloning Kit (Invitrogen). The plasmid was then used to transform LSH117 and replace the *catsacB* cassette at the *ccmA* locus with the construct. Clones that grew on HB plates containing sucrose but not chloramphenicol were selected and confirmed by PCR and Sanger sequencing (SSH38A). We noticed that these strains had altered morphology (Expanded View Figure 4) so to adjust the cell shape to be more like WT, we added a second copy of WT *ccmA* at *rdxA* by transforming LSH148 (*ccmA::catsacB, rdxA::ccmA)* with genomic DNA from SSH38A to replace *catsacB* cassette with *ccmA-40 aa linker- HaloTag* to generate strain SSH39B. Clones that grew on HB plates containing sucrose but not chloramphenicol were selected and confirmed by PCR.

To generate strains lacking *csd5, csd7,* and *csd6*, SSH39B was transformed with genomic DNA from either LSH123 (*csd5::cat*), DCY26 (*csd7::cat*), or TSH17 (*csd6::cat*). Clones that grew on HB plates supplemented with chloramphenicol were selected to generate strains SSH49A, SSH50A, and SSH70A.

To generate JS09, a strain lacking *csd5* and *csd7,* we converted the *cat* cassette to a kanamycin resistance cassette in SSH49A (*ccmA-40 aa linker-HaloTag, csd5::cat, rdxA::ccmA*) by transforming it with a PCR product containing a kanamycin resistance cassette flanked by homology to a *cat* cassette. The cassette was amplified by PCR using primers C1_828' and 'C2_829' from JTH3, then the product was purified with the QIAquick PCR Purification Kit. After transforming SSH49A with the purified PCR product we selected colonies that grew on HB plates supplemented with kanamycin but not chloramphenicol to generate strain JS02. Next, JS02 was transformed with genomic DNA from DCY28 (*csd7::cat*) and colonies that grew on HB plates supplemented with chloramphenicol and kanamycin were selected.

Construction of vectors for recombinant expression of 6-his CcmA

Truncated versions of CcmA were cloned into pET15b, a cloning vector used for expressing N-terminal 6-his tagged proteins, with the In-Fusion cloning kit (Takara). First, pET15b was linearized with restriction enzymes BamHI-HF (NEB) and NdeI (NEB), then gel extracted and purified using QIAquick Gel Extraction Kit (Qiagen). Next, PCR products containing truncated versions of CcmA flanked by 15 nucleotides of homology to the plasmid backbone were amplified and the products were purified with the QIAquick PCR Purification Kit (Qiagen).

The Δ*NT* product was amplified from SSH55A genomic DNA with primers 'CcmA-pET15b_BamH1' and 'pET15b_ccmA_delta_aa2to17_NdeI'. The Δ*MM* product was amplified from SSH54A genomic DNA with primers 'CcmA-pET15b_BamH1' and 'pET15b_ccmA_delta_aa3to12_NdeI'. The Δ*13-17* product was amplified from SSH67A genomic DNA with primers 'CcmA-pET15b_BamH1' and 'pET15b_ccmA_delta_aa13to17_NdeI'. The Δ*CT* product was amplified from SSH56B genomic DNA with primers 'CcmA-pET15b_NdeI' and 'pET15b_ccmA_delta_aa119to136_BamHI. The *BD* product was amplified from SSH57A genomic DNA with primers 'pET15b_ccmA_delta_aa2to17_NdeI' and 'pET15b_ccmA_delta_aa119to136_BamHI'.

After cloning following the manufacturer's instructions, colonies were verified by restriction digest and Sanger sequencing. Confirmed plasmids were transformed into BL21 DE3 (NEB) for recombinant expression of 6his-CcmA.

49 Spadina Ave. Suite 200
Toronto ON M5V 2J1 Canada
www.biorender.com

### Confirmation of Publication and Licensing Rights

**March 31st, 2022**
**Science Suite Inc.**

| | |
|---|---|
| ***Subscription:*** | *Institution* |
| ***Agreement number:*** | *WD23QQSNJR* |
| ***Journal name:*** | *The EMBO Journal* |

To whom this may concern,

This document is to confirm that Sophie Sichel has been granted a license to use the BioRender content, including icons, templates and other original artwork, appearing in the attached completed graphic pursuant to BioRender's Academic License Terms. This license permits BioRender content to be sublicensed for use in journal publications.

All rights and ownership of BioRender content are reserved by BioRender. All completed graphics must be accompanied by the following citation: "Created with BioRender.com".

BioRender content included in the completed graphic is not licensed for any commercial uses beyond publication in a journal. For any commercial use of this figure, users may, if allowed, recreate it in BioRender under an Industry BioRender Plan.

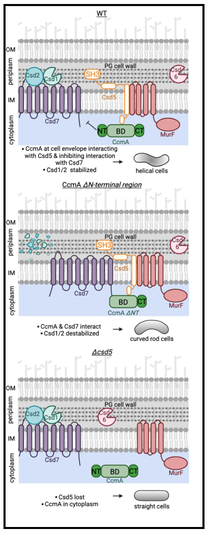

*For any questions regarding this document, or other questions about publishing with BioRender refer to our BioRender Publication Guide, or contact BioRender Support at.*
